# SARM1, the executioner of axon degeneration, is an ADP-ribosyl transferase and autoMARylation negatively regulates its activation

**DOI:** 10.1101/2025.09.12.675907

**Authors:** Janneke D. Icso, Leo DeOrsey, Leonard Barasa, Brian A. Kelch, Paul R. Thompson

## Abstract

Axon degeneration is a hallmark of nearly all neurodegenerative diseases. SARM1 plays a central role in this process by degrading NAD^+^ into nicotinamide and ADPR or cADPR. SARM1 also catalyzes a base exchange reaction between NAD^+^-phosphate (NADP^+^) and nicotinic acid (NA) to generate NAADP. These second messengers (i.e., ADPR, cADPR, and NAADP), and NAD^+^ consumption, are thought to drive axon degeneration. Herein, we identify a fourth reaction catalyzed by SARM1: mono-ADP-ribosylation (MARylation). Specifically, we show that SARM1 MARylates itself and other proteins with a catalytic efficiency (*k_cat_/K_m_*) higher than its NAD^+^ hydrolase activity. We further show that auto-MARylation promotes a phase transition and renders SARM1 responsive to regulation by NMN. Notably, endogenous SARM1 is MARylated at mitochondria, suggesting that MARylation may regulate SARM1 localization. Together, these findings uncover new regulatory mechanisms and expand the known signaling functions of SARM1.

**Significance:** SARM1 is an NAD^+^ hydrolase that executes axon degeneration in myriad neurodegenerative diseases. In addition to NAD^+^ hydrolysis, SARM1 catalyzes NAD^+^ cyclization and a base exchange reaction with NADP^+^ and nicotinic acid. These studies show that SARM1 also catalyzes the transfer of single ADPR moieties to proteins, including itself. Notably, we show that this auto-modification regulates SARM1 activity, allowing the protein to respond to NMN and to permit the phase transition. We also show that endogenous SARM1 is modified at the mitochondria, suggesting that this post-translational modification regulates SARM1 subcellular localization. These findings offer valuable mechanistic insights into SARM1 regulation that will ultimately inform the development of inhibitors targeting SARM1 for neurodegenerative diseases.

## Introduction

Wallerian degeneration is characterized by axonal fragmentation and the breakdown of the myelin sheath that encases axons.^1^ This form of axonal death closely resembles the degeneration observed in neurodegenerative diseases such as Alzheimer’s, Parkinson’s, and Amyotrophic Lateral Sclerosis (ALS), as well as in peripheral neuropathies and traumatic brain injury and is thusly called Wallerian-like degeneration.^2–7^ Sterile alpha and Toll/interleukin receptor (TIR) motif containing protein 1 (SARM1) is the central executioner of this process; SARM1 deletion or inhibition delays degeneration up to 2 weeks *in vivo* in Wallerian-like diseases.^1,8,9^ Conversely, activating mutations in SARM1 have been linked to the development of ALS.^3,10–14^ Together, these findings implicate SARM1 as a key driver of axon degeneration and underscore SARM1 as important therapeutic target for neurodegenerative diseases.^8,9^

The domain architecture of SARM1 consists of an N-terminal Armadillo repeat (ARM) domain that regulates SARM1 activity, tandem sterile alpha motif (SAM) domains that form an octameric hub, and a C-terminal TIR domain that is responsible for catalysis.^15–17^ The TIR domain hydrolyzes NAD^+^ into ADP-ribose (ADPR).^18,19^ The TIR domain catalyzes two additional reactions: the formation of cyclic-ADPR (cADPR) and a base exchange reaction between NADP^+^ and nicotinic acid (NA) to produce nicotinic acid adenine dinucleotide phosphate (NAADP).^18,20,21^ These products – ADPR, cADPR, and NAADP – are thought to act as second messengers that activate calcium channels, leading to elevated intracellular calcium levels.^22–25^ This calcium influx can then activate calpains, a class of calcium-dependent proteases,^8,9,22,26–29^ which degrade microtubules and neurofilaments, leading to axonal fragmentation and degeneration.^27,28^ Thus, SARM1 is thought to drive axon degeneration directly through NAD^+^ consumption and indirectly via calcium-mediated signaling cascades.

SARM1 activity is tightly regulated by the axonal concentration of nicotinamide mononucleotide (NMN) and a liquid-to-solid phase transition.^9,30–36^ Axonal NAD^+^ levels are controlled by nicotinamide mononucleotide adenylyltransferase 2 (NMNAT2), which synthesizes NAD^+^ from NMN and ATP.^9,37–41^ Following axonal injury, NMNAT2 levels fall, leading to a precipitous drop in NAD^+^ and a concomitant increase in the concentration of NMN.^36,37,39,42^ Under normal conditions, NAD^+^ binds an allosteric pocket in the ARM domain and maintains the enzyme in an inhibited state.^43^ Therefore, the rising NMN/NAD^+^ ratio that occurs during injury causes NAD^+^ to dissociate from the ARM domain and NMN to bind in its place.^20,35,36^ NMN binding triggers a conformational change that causes TIR domain oligomerization and a consequent increase in catalytic activity.^35,36^ SARM1 activation is further amplified by a phase transition in which SARM1 forms a supramolecular complex.^31–34^ While NMN alone increases SARM1 activity 3-4-fold, the phase transition increases SARM1 activity 23-fold and lowers the NMN activation threshold to physiologically relevant levels (i.e., ∼ 6 μM).^21,31,36,44^

SARM1 catalyzes hydrolysis, cyclization, and base exchange reactions through a shared oxocarbenium-like intermediate (**Figure 1**).^30^ This intermediate can react with small molecules such as vacor, 3-acetyl pyridine, and 5-haloisoquinolines to generate novel ADPR-adducts with diverse biological effects, including neurotoxicity, signaling, or inhibition.^20,30,45–47^ Similar chemistry is observed with plant TIR domains, which can catalyze a base exchange reaction to generate ATP-ADPR adducts.^45^ Poly-ADP-ribose polymerases (PARPs) also use an oxocarbenium-like intermediate to transfer ADPR to proteins.^48–50^ Most PARPs catalyze the transfer of only one ADPR moiety (MARylation), whereas a select few (e.g., PARP1) catalyze the transfer of multiple ADPR units (PARylation), resulting in either linear or branched polymeric chains.^49,50^ PARP activity regulates RNA stability, transcription, cell signaling and trafficking, the ubiquitin proteasome system, and the DNA damage response.^49^ Additionally, this modification can regulate phase separations, particularly in the context of stress granule formation.^51^

**Figure 1.**
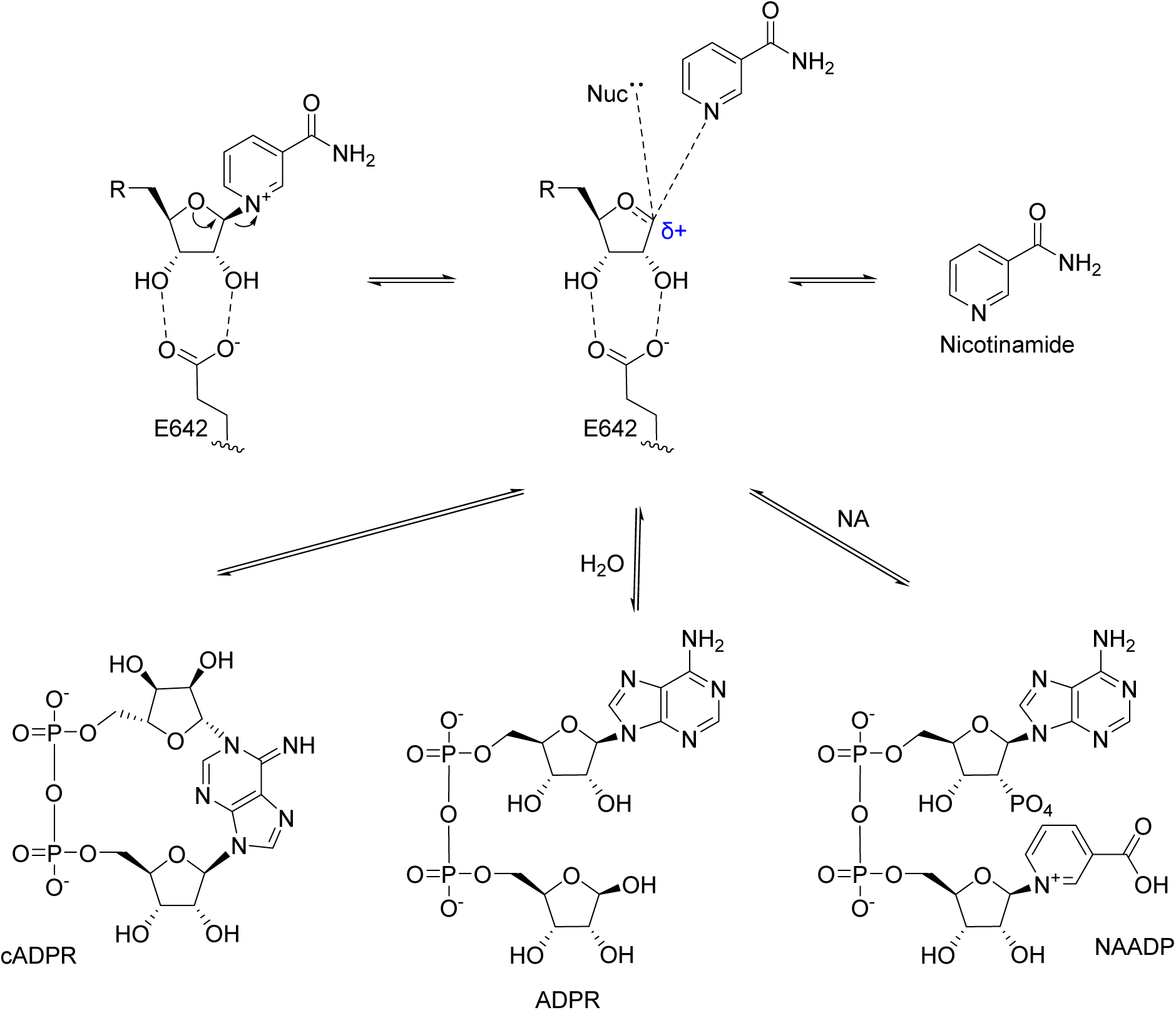
SARM1 uses an oxocarbenium-like intermediate that can also be utilized in the transfer of ADPR. The catalytic mechanism of SARM1 involves stabilization of NAD(P)^+^ by hydrogen bonding to promote nicotinamide leaving and the formation of a highly reactive oxocarbenium-like intermediate. cADPR, ADPR, and NAADP are formed by nucleophilic attack of the N1 of adenine moiety, water, or nicotinic acid (NA), respectively. R: ADPR or P-ADPR; Nuc: N1 of adenine, water, pyridine base.

Given that SARM1 and PARPs employ similar catalytic mechanisms, we hypothesized that SARM1 may also possess ADP-ribosylation activity. Indeed, we show that SARM1 exhibits PARP-like MARylation activity, modifying itself and other proteins. Overexpression of SARM1 alters cellular MARylation patterns and this activity is dependent on key catalytic residues in the TIR domain. Auto-MARylation can be reversed with hydroxylamine or MacroD1 indicating that SARM1 autoMARylates acidic residues. MARylation occurs across the protein, including domain interfaces, exposed loop regions, and adjacent to both allosteric and active sites. Moreover, we show that the removal of this PTM reduces SARM1 solubility and that MARylation is required for the enzyme to be regulated by NMN and the phase transition. Finally, we show that SARM1 catalyzes MARylation in *trans in vitro* and that endogenous SARM1 is MARylated. Together, these findings reveal a previously unrecognized layer of SARM1 regulation with broad implications for understanding axon degeneration and developing therapies for neurodegenerative diseases.

## Results

### SARM1 is an ADP-ribosyltransferase

The reactions catalyzed by SARM1 involve the formation of an oxocarbenium-like intermediate (**Figure 1**). Given that PARPs use a similar catalytic mechanism, we hypothesized that SARM1 might possess a fourth enzymatic activity and catalyze the transfer of ADP-ribose to proteins. To initially validate this hypothesis, we expressed a Protein A-tagged SARM1 construct lacking the mitochondrial localization sequence (MLS; SARM1^ΔMLS^) in Expi293F cells.^30,31^ The MLS is dispensable for the catalytic activity of SARM1 and its role in axon degeneration, but its removal improves protein solubility.^17,30^ We lysed cells by sonication and evaluated the ADP-ribosylation state of proteins in the lysate by Western blot using antibodies that recognize MARylated or PARylated proteins.^52^ The ribosylation states of several proteins increased when SARM1 is overexpressed in Expi293F cells, which do not express SARM1 endogenously. For example, only a single 130 kDa protein is MARylated in the untransfected control. By contrast, in the presence of SARM1, multiple proteins are MARylated, including bands at 25, 40, 80, and 200 kDa. Notably, a band corresponding to the size of the SARM1^ΔMLS^-Protein A construct (i.e. 77 kDa) is both MAR- and PARylated (**Figures 2A, S1A**).

**Figure 2.**
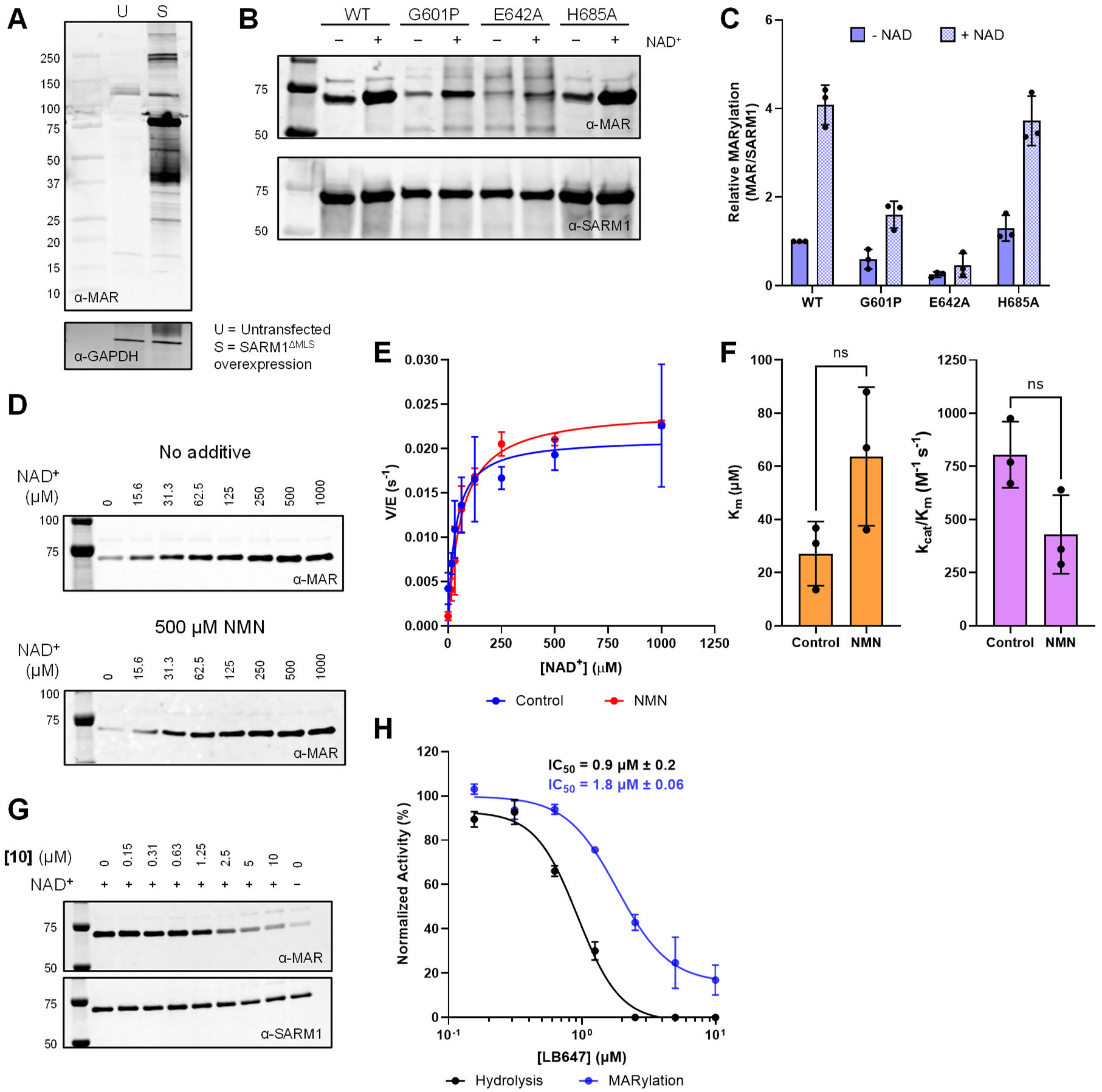
SARM1 has ADP-ribosylation activity *in vitro*. A) Recombinantly expressed SARM1 is mono-ADP-ribosylated in Expi293F lysates. U: untransfected; S: SARM1^ΔMLS^ overexpression. n = 3, representative image shown. B) Purified recombinant SARM1 has MARylation activity that is dependent on the catalytic residue. Purified WT or variant SARM1^ΔMLS^ incubated with or without 1 mM NAD^+^ for 30 min and analyzed by Western blot. n = 3, representative image shown. C) Quantification of B. Error reported as SD. D) Kinetics of MARylation *in vitro*. 750 nM SARM1^ΔMLS^ incubated with a 2-fold serial dilution of 0-1000 μM NAD^+^ in the presence or absence of 500 μM NMN and analyzed by Western blot. n = 3, representative image shown. E) Michaelis-Menten plots from D. Error reported as SD. F) The Michaelis constant (*K_m_*) and catalytic efficiency (*k_cat_/K_m_*) from D and E. Error reported as SD. G) Inhibition of the MARylation reaction by compound **(10),** a known SARM1 inhibitor. SARM1^ΔMLS^ was incubated with 0-10 μM **(10)** and then with 1 mM NAD^+^ and analyzed by Western blot. n = 3, representative image shown. H) Quantification of G. Error reported as SD. Note: in some cases, the error is smaller than the size of the data point and is not visible.

To confirm that SARM1 has intrinsic ADP-ribosylation activity, as opposed to an indirect effect caused by its overexpression, we purified SARM1^ΔMLS^ by IgG affinity purification and cleaved off the Protein A tag. SARM1^ΔMLS^ was incubated with and without NAD^+^ and the ribosylation state of the purified protein was analyzed by Western blotting. SARM1^ΔMLS^ was MARylated at baseline, and the MARylation level increased 4-fold in the presence of NAD^+^ (**Figure 2B and C**). These data indicate that SARM1 is ribosylated in cells and that SARM1 can modify itself *in vitro*.

Having demonstrated that SARM1 autoMARylates, we next sought to confirm that this activity depends on its catalytic function and/or oligomerization status. For these studies, we generated mutants designed to disrupt either enzymatic activity (i.e., the E642A mutant) or oligomerization (i.e., the G601P and H685A mutants). E642 interacts with the hydroxyl groups present on the ribose adjacent to the nicotinamide moiety and these interactions are important for stabilizing the oxocarbenium-like intermediate (**Figure 1**).^30,46,47^ By contrast, G601 and H685 are located at key multimerization interfaces and their mutation disrupts oligomerization.^16,32,33^

In the absence of exogenous NAD⁺, the basal MARylation levels of the E642A and G601P mutants was minimal, i.e. 2-3-fold less than the wildtype enzyme. By contrast, the H685A mutant showed MARylation levels comparable to wild-type SARM1^ΔMLS^. Upon NAD⁺ supplementation, the H685A mutant maintained wild-type activity, while the G601P mutant displayed approximately 50% of the MARylation observed in the wild-type. Notably, the E642A catalytic mutant showed a 3-fold reduction in MARylation even in the presence of NAD⁺ (**Figure 2B and C**). These results suggest that MARylation by SARM1^ΔMLS^ is dependent on its catalytic activity but largely independent of its oligomerization state.

Surprisingly, we did not observe significant PARylation of purified wild type SARM1^ΔMLS^ or several mutants that lack catalytic activity or the ability to oligomerize (**Figure S1A-C**). To reconcile these results, we added pure SARM1^ΔMLS^ to untransfected HEK293T cell lysates and evaluated the effect on protein PARylation. Notably, there was no increase in PARylation. The same pattern was apparent for the catalytically impaired E642A mutant (**Figure S1D**). We do, however, observe consistent PARylation of the SARM1^ΔMLS^-Protein A construct (**Figure S1D**). Consequently, we conclude that the Protein A tag is PARylated when overexpressed.

Given that G601 and H685 are critical for the NAD^+^ hydrolysis activity of SARM1 in the context of the isolated TIR domain,^32,33,53^ it was surprising that these oligomerization mutants retain their MARylation activity. Moreover, the G601P mutant delays axon degeneration by 5-fold in axotomy models.^16^ Therefore, we determined the kinetic parameters for the NAD^+^ hydrolysis reaction for both the G601P and H685A mutants using wild type SARM1^ΔMLS^ and the E642A mutant as controls. In buffer alone, G601P had 100-fold decreased activity compared to wild type SARM1. E642A and H685A had no detectable activity (**Figure S1E and F**). Incubating SARM1 with molecular crowding agents like PEG 3350 causes the enzyme to undergo a phase transition that activates the enzyme.^9,30–33^ NMN binds an allosteric pocket in the ARM domain to activate the enzyme.^21,35,36,39^ Notably, the combination of the phase transition and NMN increase SARM1 activity 23-fold.^31^ Under conditions in which the enzyme is maximally activated (i.e., 25% PEG 3350 and 500 mM NMN), the H685A mutant had a 7.5-fold reduction in activity, whereas the G601P and E642A mutants had no detectable activity (**Figure S1E and F**). Therefore, oligomerization is required for NAD^+^ hydrolysis but not MARylation.

To characterize the kinetics of the MARylation reaction, we performed end-point kinetic experiments where SARM1 was incubated with 0-1000 µM NAD^+^ for 1 min and changes in MARylation monitored by Western blotting. The *K_m_* for the MARylation reaction was approximately 30 µM (**Figure 2D-F, S1G and H**), which is consistent with the *K_m_* for the NAD^+^ hydrolysis reaction and 6-fold less than ENAD hydrolysis reaction.^20,31^ By contrast, the catalytic efficiency (*k_cat_/K_m_*) of the MARylation reaction was 750 M^-1^s^-1^ (**Figure 2D-F**), which is 7.5-fold higher than either the NAD^+^ or ENAD hydrolysis reactions,^20,31^ suggesting that the MARylation reaction is physiologically relevant.

To complete our initial characterization of the autoMARylation activity of SARM1, we evaluated whether inhibitors of the hydrolysis reaction also block autoMARylation. Increasing concentrations of an isothiazole-based inhibitor, compound **10**,^54^ were incubated with SARM1 for 30 min and changes in autoMARylation measured by Western blotting. Compound **10** dose-dependently inhibits the autoMARylation activity of SARM1 with an IC_50_ of 1.8 µM. This value is similar to the one obtained for the NAD^+^ hydrolysis reaction (IC_50_ = 930 nM, **Figure 2G and H**). Since **10** is a covalent inhibitor, the ∼2-fold difference in the IC_50_ values likely reflects the different incubation times used rather than a meaningful difference in the inhibition of one reaction over the other. We also tested zinc chloride, another SARM1 inhibitor,^32,55^ and found that the IC_50_ for the autoMARylation reaction is 2.1 µM which is comparable to the IC_50_ for the hydrolysis reaction (IC_50_ = 1.1 µM, **Figure S1G and H**). Considered together, these data indicate that SARM1 uses a common intermediate to catalyze autoMARylation, NAD(P)^+^ hydrolysis, cyclization, and base exchange.

### MARylation inhibits both the activity and activation of SARM1

MARylation can occur on Asp, Glu, Cys, Arg, Lys and Ser residues. To ascertain what residues are modified during the autoMARylation of SARM1, we first examined whether this PTM could be removed with hydroxylamine (NH_2_OH); NH_2_OH transacylates the modified protein to remove the ADP-ribose modification from aspartate and glutamate residues, leaving a hydroxamic acid derivative in place of ADPR (**Figure 3A**).^56^ For these studies, SARM1^ΔMLS^ was incubated with and without NAD^+^ for 20 min, with and without NH_2_OH for 1 h, and then the samples were analyzed by Western blotting. In the presence of NAD^+^, SARM1^ΔMLS^ MARylation increased from undetectable levels. By contrast, MARylation was absent in samples treated with both NAD^+^ and then NH_2_OH (**Figure 3B**). These data indicate that NH_2_OH effectively replaces ADP-ribosylation on SARM1^ΔMLS^ and that SARM1 MARylation occurs primarily on acidic residues. Next, we determined whether MacroD1 could reverse the autoMARylation activity of SARM1. MacroD1 is an enzyme that removes single ADPR moieties from proteins, particularly from aspartate and glutamate residues.^50^ In contrast to NH_2_OH, MacroD1 does not leave a scar upon ADPR removal, thus yielding the unmodified protein (**Figure 3B**). For these studies, we first incubated SARM1^ΔMLS^ with NAD^+^ for 5 min to autoMARylate SARM1, and then added MacroD1 for 15, 30, or 60 min. Similar to the results obtained with NH_2_OH, MARylation decreased to basal levels within 15 min and remained at baseline at 1 h (**Figure 3D**). These data confirm that SARM1 is MARylated at aspartate and glutamate residues.

**Figure 3.**
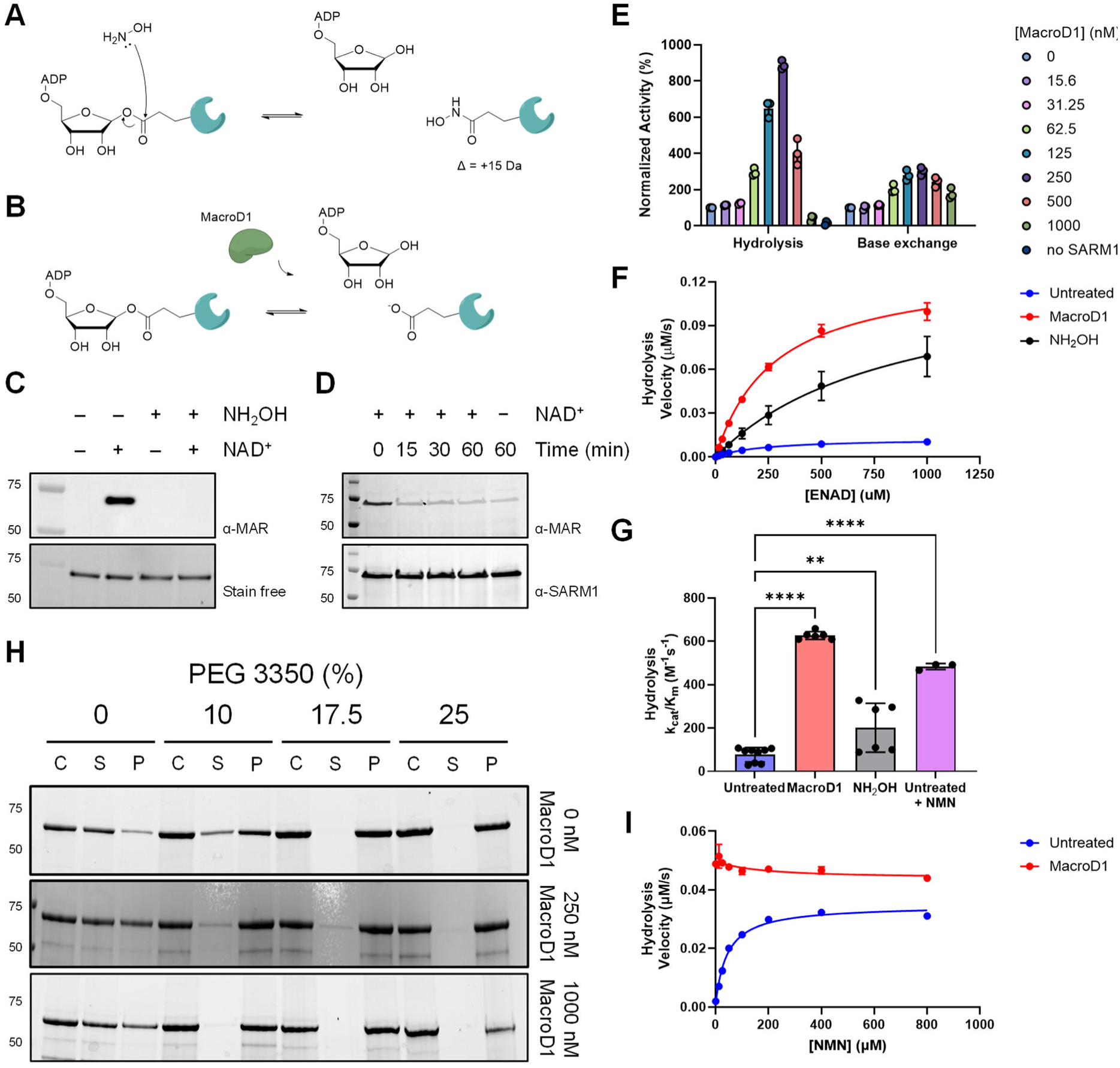
MARylation inhibits the activity and activation of SARM1. A) Mechanism by which hydroxylamine removes ADP-ribosylation from proteins. B) Mechanism by which MacroD1 removes ADPR moieties from proteins to yield unmodified residues. C) Hydroxylamine removes MARylation from SARM1; n = 3, representative image shown. D) MacroD1 removes MARylation from SARM1 over 1 h; n = 2, representative image shown. E) MacroD1 dose response; n = 3, error reported as SD. F) SARM1^ΔMLS^ ENAD hydrolysis activity in the presence and absence of 500 mM hydroxylamine or 250 nM MacroD1; minimum n = 3, error reported as SD. G) Catalytic efficiency of ENAD hydrolysis in the presence and absence of 500 mM hydroxylamine or 250 nM MacroD1 from F. Data for the Untreated + NMN condition was extrapolated from Icso et al. (2023). H) Coomassie stained SDS-PAGE gels of centrifugation-based phase transition experiments in PEG 3350; n = 2, representative image shown. I) NMN dose response in the hydrolysis reaction of SARM1^ΔMLS^ treated with and without 250 nM MacroD1; n = 3, error reported as SD. Note: in some cases, the error is smaller than the size of the data point and is not visible.

Having demonstrated that SARM1 autoMARylates at aspartate and glutamate residues, we next sought to determine whether MARylation affects the activity of SARM1. Since SARM1 is partially autoMARylated during expression, we first sought to determine how the removal of this PTM would affect enzymatic activity. Here, SARM1^ΔMLS^ was first incubated with increasing concentrations of MacroD1 (0-1 µM) and then the hydrolysis and base exchange activities were analyzed. Hydrolysis activity was measured using an NAD^+^ analog, etheno-NAD^+^, that fluoresces upon nicotinamide cleavage.^33,57^ Base exchange activity was assayed using pyridyl conjugate 6 (PC6) which generates a fluorescent product, i.e. PC6-adenine dinucleotide (PAD6), in the presence of NAD^+^.^30,31,58^ Notably, both the hydrolysis and base exchange activities showed parabolic dose-response curves, where the activities were optimal at 250 nM MacroD1 (**Figure 3E**). With respect to the hydrolysis reaction, enzymatic activity was 9-fold greater at 250 nM MacroD1 than in the absence of MacroD1. Base exchange activity was 3-fold greater under the same conditions. By contrast, treatment with 1 µM of MacroD1 led to a 20-fold reduction in the hydrolysis activity (relative to the rate at 250 nM MacroD1); the base exchange activity was 2- fold lower (**Figure 3E**). These data suggest that autoMARylation controls SARM1 activity, both negatively and positively, depending on the extent of this PTM.

To investigate this hypothesis further, we examined the hydrolysis activity of SARM1^ΔMLS^ in the presence and absence of 500 mM NH_2_OH or 250 nM MacroD1. In the presence of NH_2_OH, the catalytic efficiency (*k_cat_/K_m_*) for ENAD hydrolysis was 2-fold greater than the control and 6- fold higher in the presence of MacroD1 (**Figure 3F and G**). For comparison, NMN increases the hydrolysis activity of SARM1 by 5-fold (**Figure 3G**).^31,35,36^ Thus, MacroD1 treatment and NMN modulate SARM1 activity to a similar extent. The smaller effect observed with NH_2_OH is likely attributable to the fact that NH_2_OH leaves a neutral hydroxamic acid scar, which could cause conformational changes that minimize the impact of removing a MARylation modification.

Since the phase transition and NMN activate SARM1,^31^ we further hypothesized that MARylation might impact the effect of these two regulation mechanisms. To test this hypothesis, we incubated SARM1^ΔMLS^ with 25% PEG 3350 in the presence and absence of 500 µM NMN and measured the hydrolysis activity. Consistent with our previous data, the combination of PEG 3350 and NMN enhanced SARM1 activity by 25-fold. When 250 nM MacroD1 was incubated with SARM1, the hydrolysis activity increased to a similar extent. However, PEG and NMN did not significantly alter the effect of MacroD1 (**Figure S2A**). These data indicate that MARylation negatively regulates SARM1 activity independently of PEG and NMN.

Next, we sought to determine whether MacroD1 impacts the phase transition. Briefly, SARM1^ΔMLS^ was incubated first with 250 or 1000 nM MacroD1 for 30 min and then with 0-25% PEG 3350 for another 15 min. Thereafter, the samples were centrifuged at 21,000 *xg* and the supernatant and pelleted fractions were separated and analyzed by SDS-PAGE. In the absence of MacroD1, SARM1^ΔMLS^ was primarily located in the supernatant fraction at 0% PEG 3350. By contrast, at 10-25% PEG 3350, SARM1^ΔMLS^ was found in the pellet (**Figure 3H**), consistent with our previous results.^30–33^ In the presence of MacroD1, SARM1^ΔMLS^ was observed in both the supernatant and pellet fractions without PEG, whereas the protein was increasingly located in the pelleted fraction with PEG 3350. Notably, this pattern is dose dependent on MacroD1, where SARM1 is predominantly located in the pellet fraction at lower PEG concentration at 1000 nM MacroD1 (**Figure 3H**). These results indicate that MARylation improves SARM1 solubility and partially blocks the phase transition in the absence of molecular crowding agents.

Finally, we examined the effect of MARylation on the NMN response. For these studies, SARM1^ΔMLS^ was incubated with and without MacroD1, NMN (0-800 µM) was then added, and SARM1 activity evaluated in the hydrolysis and base exchange assays. For the hydrolysis reaction, the EC_50_ of NMN is 40 µM in the absence of MacroD1. By contrast, when SARM1^ΔMLS^ is pretreated with MacroD1, the enzyme is already fully activated in the absence of NMN, i.e. SARM1 activity is independent of NMN concentration (**Figure 3I**). Similar results were observed in the base exchange assay (**Figure S2C**). Overall, these data indicate that MARylation regulates SARM1 activation by affecting both the solubility of SARM1 and its ability to respond to NMN.

### MARylation of the SS loop permits SARM1 activity through the phase transition

In total, our data indicates that SARM1 has an optimal level of MARylation. To identify the residues mediating these effects, we employed mass spectrometry. Although mono-ADP- ribosylation (MARylation) results in a substantial +541 Da mass shift, the modification is labile and difficult to detect using conventional methods. To address this challenge, hydroxylamine (NH₂OH) is commonly used to facilitate site identification, as it leaves a stable hydroxamic acid “scar” on the protein, resulting in a +15 Da mass shift that is more amenable to mass spectrometric analysis. For our studies, SARM1^ΔMLS^ was incubated with and without NAD^+^ for 20 min at room temperature to induce auto-MARylation and then treated with 500 mM NH_2_OH for 1 h to promote the transacylation reaction. Thereafter, proteins were processed for tandem mass spectrometry. In total, 20 modification sites were identified across all domains in SARM1 (**Figure 4A-D**). In the ARM domain, modified residues included D127, E155, E189, and D367. Interestingly, E189 is located adjacent to the allosteric pocket in the ARM domain that binds NAD^+^ and NMN (**Figure 4A and B**). In the SAM domains, the modification sites line the center of the octameric ring (i.e., 450, 454, 520, 525, and 526) or are located at domain interfaces (i.e., 414, 416, and 489, **Figure 4A**). In the TIR domain, key modification sites included D594, which is adjacent to the active site, and two residues in the SS loop, D632 and D634. Additional modified residues in the TIR domain included D596, E686, E689, and E693 (**Figure 4A and B**).

**Figure 4.**
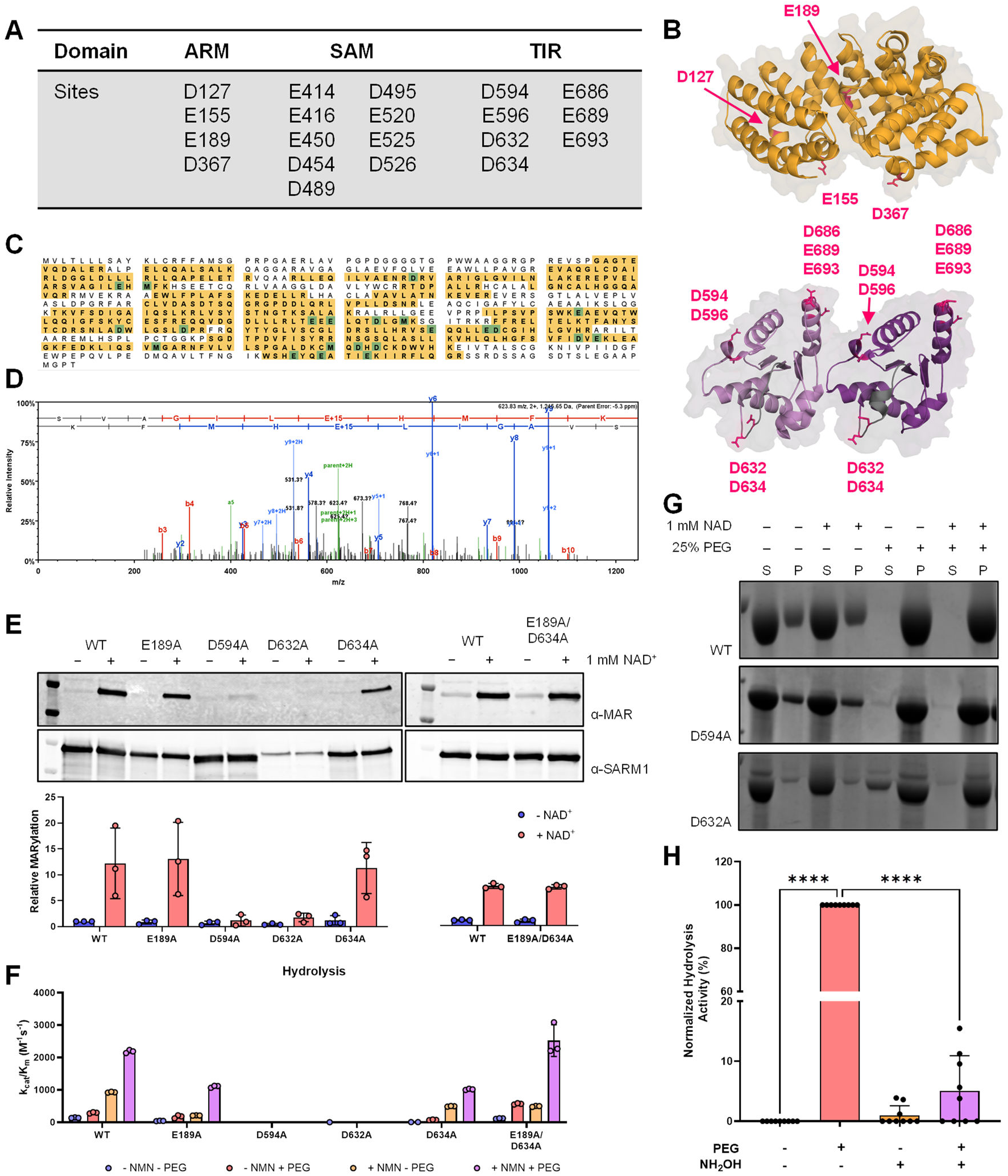
MARylation of the TIR domain is required for activity. A) Summary of ribosylation sites across the full-length protein. B) Modification sites mapped to ARM (top, PDB: ) and TIR (bottom, PDB: ) domain structures. Sites of modification are highlighted in pink. C) Coverage of SARM1 (60%) from mass spectrometry experiments used to identify the sites of modification. D) Representative MS2 spectra. E) MARylation of SARM1^ΔMLS^ variants. Western blot for MARylation and SARM1 (top). Relative quantification of blots (bottom). n = 3; representative images shown and error reported as SD. F) Catalytic efficiency with respect to the hydrolysis reaction of MARylation variants treated with 0-1000 µM ENAD. n = 3, error reported as SD. G) Coomassie stained SDS-PAGE for centrifugation-based phase transition experiments of the SARM1 MARylation variants. n = 3, representative images shown. H) Hydrolysis activity of the TIR domain from SARM1 treated with and without 25% PEG 3550 or 500 mM hydroxylamine. n = 3, error reported as SD. Note: in some cases, the error is smaller than the size of the data point and is not visible.

We selected E189, D594, D632, and D634 for further analysis because these residues are in regulatory regions or adjacent to the active site. First, we generated the E189A, D594A, D632A, D634A, and E189A/D634A mutants and evaluated their ability to autoMARylate. For these studies, wild type and mutant SARM1^ΔMLS^ were incubated with 1 mM NAD^+^ for 15 min and MARylation levels were evaluated by Western blot. MARylation of wild type SARM1 increased ∼10-fold in the presence of NAD^+^. The E189A, D634, and E189A/D634A variants were MARylated like wild type SARM1^ΔMLS^, also displaying 10-fold increases MARylation after NAD^+^ incubation (**Figure 4E**). These results indicate that E189A and D634 are not primary MARylation sites. By contrast, we observed little MARylation of the D594A and D632A variants (**Figure 4E**).

Since the lack of MARylation could be due to an effect on enzymatic activity (rather than the removal of a site of ADP-ribosylation), we evaluated the hydrolysis and base exchange activity of all the mutants (**Figure 4F and S3**). The D594A and D632A variants lacked any detectable catalytic activity. The lack of activity may relate to the proximity of D594 and D632 to the active site which impairs both the MARylation and hydrolysis activities of SARM1.

Consistent with their ability to autoMARylate, the E189A, D634A, and E189A/D634A mutants were all active. Specifically, the activity of the E189A and D634A mutants were decreased by 2-fold compared to wild-type, whereas the E189A/D634A double mutant had near wild type-like activity at baseline. Notably, the D634A mutant could be progressively activated by PEG, NMN, or a combination of PEG and NMN to a maximum of half wild-type activity under full activation conditions (i.e., 25% PEG and 500 µM NMN). E189A and E189A/D634A respond to activation by PEG or a combination of NMN and PEG, but not NMN alone. Specifically, E189A and E189A/D634A showed 2.5-fold increased hydrolysis activity in PEG or NMN whereas wild type activity increased 2.5-fold and 4-fold in 25% PEG or 500 µM NMN, respectively (**Figure 4F and S3**). These results indicate that the NMN dependence of the E189A and E189A/D634A variants is impaired in the absence of the phase transition and suggest that MARylation at E189 is an additional pathway to activate SARM1, though SARM1 activation via E189 MARyation is likely minor.

We also measured the effect of mutating these residues on the base exchange activity of SARM1. Notably, the base exchange activity of the E189A and D634A mutants was significantly impaired (8-fold) in the absence of additives. By contrast, the base exchange activity of the E189A/D634A variant was increased by 2-fold. In the presence of PEG 3350, the base exchange activity of wild type SARM1 is significantly inhibited overall. The activity of the E189A, D634A, and E189A/D634A mutants was similarly impaired under the same conditions. In the presence of NMN, the base exchange activities of the E189A, D634A, and E189A/D634A variants to the level of wild type SARM1^ΔMLS^ (**Figure S3**). Our fluorescent base exchange assay uses pyridyl conjugate 6, which has a high apparent affinity for SARM1 in the absence of PEG and we believe these data reflect this affinity rather than any real regulation of the product specificity of these MARylation mutants.^30^

Because MARylation promotes solubility and prevents the phase transition in the absence of additives (**Figure 3H**), we sought to determine how the removal of a MARylation site would impact the phase transition. As such, SARM1^ΔMLS^ mutants were incubated in the presence and absence of 25% PEG 3350 for 15 min and centrifuged at 21,000 *x g.* The supernatant and pelleted fractions were separated and analyzed by SDS-PAGE. In the absence of PEG, wild type SARM1 was located primarily in the supernatant, whereas the protein was in the pellet in the presence of PEG. Pre-incubation with 1 mM NAD^+^ to MARylate the protein did not affect this pattern (**Figure 4G**). The D594A variant transitioned like wild type SARM1. By contrast, D632A was found in the supernatant in the absence of PEG and split between the supernatant and pellet fractions in the presence of PEG (**Figure 4G**). The D632A mutant preincubated with NAD^+^ was found in the pellet in the presence of PEG, indicating that this mutant undergoes the phase transition under these conditions. However, this result is not due to autoMARylation because this mutant does not have MARylation activity (**Figure 2B and C**). These data indicate D632A variant does not completely undergo the phase transition and suggest that MARylation at the SS loop is critical for TIR domain activity.

We hypothesized that MARylation of the TIR domain might affect the ability of PEG to activate the catalytic activity of this domain. To test this hypothesis, the TIR domain from SARM1 was treated with NH_2_OH and then incubated in the presence and absence of PEG 3350 to initiate the phase transition. When we tested the activity of the TIR domain in the hydrolysis assay, we observed strong activity only in the presence of PEG. Consistent with our hypothesis, NH_2_OH decreases hydrolysis activity by 20-fold (**Figure 4H**). Considered together, these results suggest that the TIR domain, specifically the SS loop, is MARylated to permit the phase transition.

### MARylation occurs in trans

To further characterize the autoMARylation activity of SARM1, we sought to determine whether this modification occurs in *cis* (i.e., the same protein modifies itself) or in *trans* (i.e., one protein monomer modifies another protein). To distinguish between these two possibilities, we incubated the TIR domain (hTIR) with a catalytically inactive SARM1^ΔMLS^ variant (E642A) and examined the MARylation levels by Western blotting. Purified hTIR in the absence of NAD^+^ is MARylated (**Figure 5A**). In the presence of NAD^+^, MARylation of hTIR increases >10-fold (**Figure 5A**). By contrast, the E642A SARM1^ΔMLS^ variant showed minimal MARylation in the presence and absence of NAD^+^. When the E642A variant was incubated with increasing concentrations of hTIR, we observed a dose-dependent increase in MARylation of the E642A construct, indicating that hTIR can MARylate a distinct substrate and that MARylation can occur in *trans* between separate SARM1 monomers (**Figure 5A**). Having shown that auto- MARylation occurs in *trans*, we next determined whether SARM1 modifies other proteins. Specifically, we repeated the above experiment but substituted the E642A SARM1^ΔMLS^ mutant with protein arginine deiminase 2 (PAD2), another protein commonly used in our laboratory. Similar to E642A, we observed a dose-dependent increase in PAD2 MARylation with TIR domain concentration (**Figure 5B**), indicating that SARM1 can MARylate non-SARM1 substrates.

**Figure 5.**
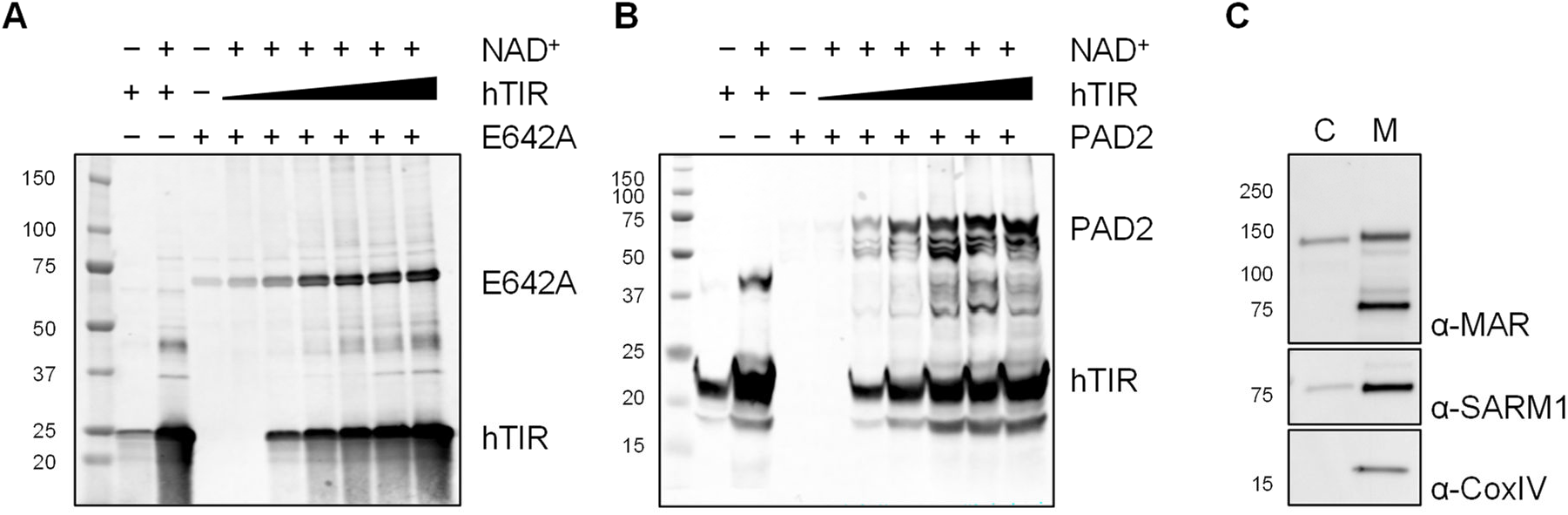
SARM1 catalyzes MARylation in *trans*. A) SARM1^E642A^ MARylation by the TIR domain *in vitro*. 750 nM E642A, 2.5 uM hTIR, and 0-4 uM hTIR incubated with and without 1 mM NAD^+^. n = 2, representative image shown. B) PAD2 MARylation by the TIR domain *in vitro*. 750 nM PAD2, 2.5 uM hTIR, and 0-4 uM hTIR incubated with and without 1 mM NAD^+^. n = 2, representative image shown. C) Western blot for SARM1, MARylation, and CoxIV from SHSY-5Y cells (C = cytosolic, M = mitochondrial). n =3; representative images shown.

### Endogenous SARM1 autoMARylates

Having demonstrated that overexpressed and purified SARM1 can autoMARylate, we next sought to determine whether endogenously expressed SARM1 can autoMARylate. For these experiments, we performed subcellular fractionation methods in the neuron-like cell line SHSY5Y to enrich for mitochondria. Following a 4 h incubation in serum-free media, mitochondria were isolated and analyzed by Western blot for SARM1 localization and proteome-wide MARylation. The results demonstrated that in SHSY5Y cells, SARM1 localizes to both the cytosol and mitochondria, though 10-fold more extensively to the mitochondria. Notably, autoMARylated SARM1 was observed exclusively at the mitochondria (**Figure 5C**). These results suggest that MARylation of endogenous SARM1 regulates its localization.

Lastly, we hypothesized that SARM1 also catalyzes the MARylation of other proteins in cells because SARM1 modifies PAD2 *in vitro*. To test this hypothesis, we overexpressed tag-free and Protein A-tagged SARM1^ΔMLS^ in Expi293F cells, performed subcellular fractionation, and analyzed MARylation by Western blotting. Overexpression of tag-free SARM1^ΔMLS^ did not appear to affect proteome-wide MARylation, though the expression of tag-free SARM1^ΔMLS^ was lower than Protein A-tagged SARM1^ΔMLS^. By contrast, proteome-wide MARylation increased dramatically with Protein A-tagged SARM1 overexpression and was slightly greater in the cytosolic fraction. Notably, Protein A-tagged SARM1^ΔMLS^ appeared equally split between the cytosolic and mitochondrial fractions despite the absence of the MLS, suggesting that mitochondrial localization of SARM1 is dependent on protein-protein interactions. Regardless of localization, Protein A-tagged SARM1^ΔMLS^ was MARylated in cells (**Figure S4**).

## Discussion

SARM1 catalyzes the hydrolysis and cyclization of NAD^+^ and a base exchange reaction between NADP^+^ and NA.^20,21,30,59^ The products of these reactions contribute to an influx of calcium into the axon, ultimately resulting in the axonal fragmentation characteristic of Wallerian-like diseases.^27,29^ These different reactions all proceed through the same oxocarbenium intermediate.^30^ Importantly, the oxocarbenium intermediate is also the chemical foundation for ADP-ribosylation reactions. In fact, PARP and SARM1 employ similar catalytic mechanisms. Our data demonstrate that SARM1 is also an ADP-ribosyltransferase that MARylates itself up to 10- fold when incubated with NAD^+^ (**Figure 4E**). This activity is dependent on the catalytic activity of SARM1 as the E642A mutant does not autoMARylate. Although oligomerization is required for the NAD^+^ hydrolase activity of SARM1, it is not required for MARylation because the G601P and H685A oligomerization mutants maintain their ability to autoMARylate (**Figure 2B and C**). The auto-modification occurs across the entire protein with notable sites of modification adjacent to the allosteric and active sites and in the SS loop (**Figure 4A and B**). AutoMARylation of most of these sites appears to be substochiometric as we do not observe a significant mass shift in the protein when analyzed by Western blotting. Nevertheless, it is notable that the catalytic efficiency of the MARylation reaction is 7.5-fold greater than the hydrolysis reaction (**Figure 2H**), indicating that a portion of NAD^+^ catalysis by SARM1 contributes to its auto-modification.

PTMs represent a major mechanism by which signaling pathways and individual proteins are regulated. PTMs can alter the localization, activation, solubility, functionality, and stability of proteins. ADP-ribosylation specifically is associated with many signaling events including, but not limited to, alterations in protein-protein interactions, phase separations, and macromolecular stability.^49^ For example, ADP-ribosylation is important for stress granule formation, the recruitment of ubiquitin ligases, and transcription initiation.^49,50,60^ Notably, SARM1 is ribosylated in a neuronal cell line (e.g., SHSY5Y cells, **Figure 5C**), and our data suggest that autoMARylated SARM1 preferentially localizes to mitochondria.

SARM1 activity is known to be regulated by the levels of NMN and a phase transition. Importantly, we show that MARylation modulates both regulators. For example, we showed that the catalytic activity of SARM1 is dependent on its MARylation level. Higher levels of MARylation correlate with less activity, moderate levels with optimal activity, and low levels with less activity (**Figure 3G**). These data indicate that the MARylation of specific residues differentially regulates SARM1. For example, D632 hydrogen bonds with the backbone of other residues in the SS loop and mutating this residue to alanine eliminates a hydrogen bond that stabilizes the conformation of the SS loop. Without hydrogen bond acceptors from D632 itself or from an ADPR moiety at this residue after MARylation, the protein does not undergo the phase transition (**Figure 4G**). Similarly, E189A is located in the allosteric site in the ARM domain that binds NMN. Mutagenesis at this residue unsurprisingly eliminates the response to NMN in buffer alone. However, E189A variants are NMN responsive in the phase transitioned state (**Figure 5A**). Therefore, if MARylation of E189 occurs before NMN binding, the modification likely locks the ARM domain in an active conformation, similar to the binding of NMN. These data highlight auto- MARylation as an additional regulatory mechanism of SARM1.

Together, these findings highlight autoMARylation as an additional regulatory mechanism for SARM1 and support a model in which SARM1 is MARylated at baseline to inhibit SARM1 activity and then SARM1 is further MARylated to stabilize SARM1 activation (**Figure 3H, 4G and H**). Specifically, MARylation is required for the enzyme to be activated by NMN and MARylation inhibits the phase transition (**Figure 3H and I**). After undergoing the phase transition, the levels of NMN needed to fully activate SARM1 are reduced and the subsequent increase in MARylation stabilizes the phase transition (**Figure 4G**, **Figure 6**). In addition to these effects, the autoMARylation activity of SARM1 may alter protein-protein interactions and consequently which substrates are modified. Ultimately, the data presented above describe a novel, fourth reaction type catalyzed by SARM1 and provide important information about the regulation of SARM1.

**Figure 6.**
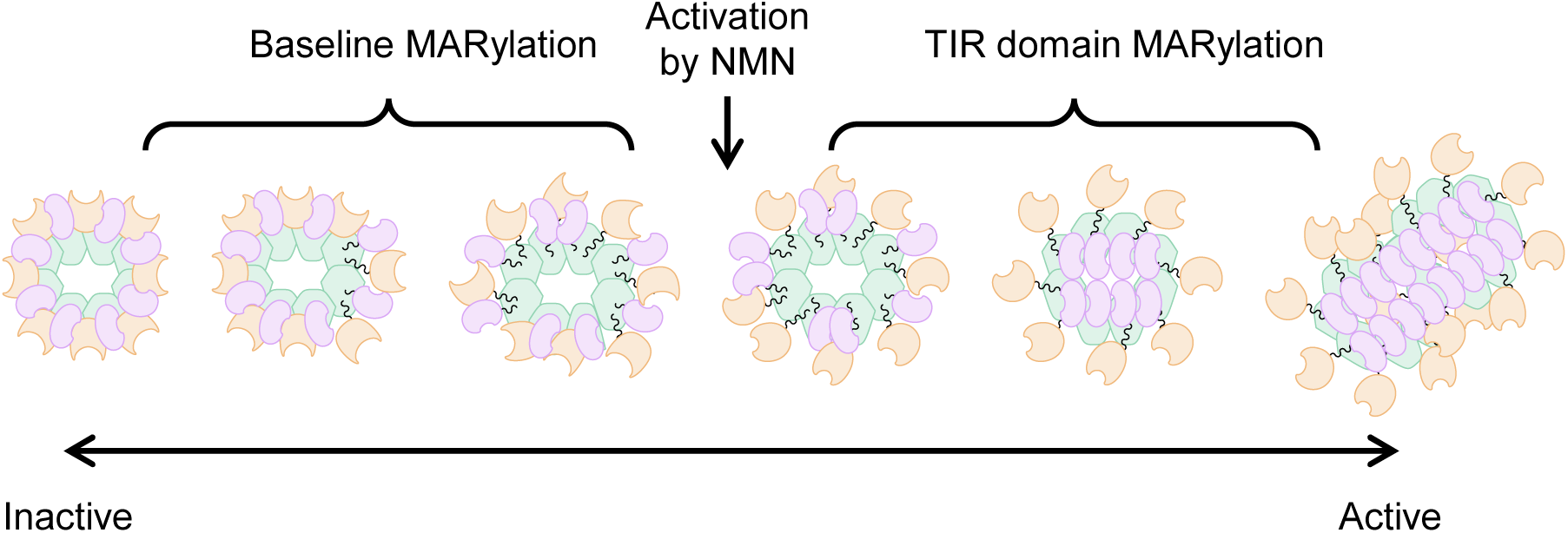
Model for SARM1 regulation by auto-MARylation.

## Materials and Methods

### SARM1^ΔMLS^ expression and purification

The expression and purification of SARM1^ΔMLS^ was performed as previously described.^30,31^ Briefly, a pcDNA3.4(+) expression construct encoding amino acids 28-724 of SARM1, a PreScission Protease site, and two tandem Protein A tags was transfected into Expi293F cells. 2.5-3 × 10^6^ cells/mL were seeded in FreeStyle 293 Expression Medium (Gibco) and incubated for 18 h at 37 °C, 8% CO_2_, < 80% relative humidity, and 125 rpm agitation. The next day, cells were diluted to 2.5 × 10^6^ cells/mL in 30 mL of culture. 3 μg plasmid DNA per mL of transfection volume was diluted in 1.5 mL OptiMEM (Gibco). 3 μg PEI MAX (PolyScience) per 1 μg DNA was diluted in 1.5 mL OptiMEM. DNA and PEI MAX dilutions were incubated individually for 5 min at room temperature after which the dilutions were mixed 1:1 and incubated 30 min further at room temperature. Dropwise, the DNA:PEI MAX complexes were added to the cultures. Transfection occurred overnight at 37 °C, 8% CO_2_, < 80% relative humidity, and 125 rpm agitation. The following day, valproic acid was dissolved in Freestyle media such that the concentration was 4.4 mM. 30 mL of valproic acid supplemented media was added to 30 mL of culture so that the final concentration of valproic acid was 2.2 mM. SARM1^ΔMLS^ was allowed to express for 3-4 days at 37 °C, 8% CO_2_, < 80% relative humidity, and 125 rpm agitation. Cells were collected by centrifugation and the media as removed by aspiration. Cell pellets were flash frozen in liquid nitrogen and stored at -80 °C until use.

To purify SARM1^ΔMLS^, cells from a total of 300 mL of culture were thawed on ice. Cells were resuspended in 10 mL lysis/wash 1 buffer (50 mM HEPES, pH 8.0; 400 mM NaCl; 5% glycerol [w/v]; Pierce Protease Inhibitor Mini Tablets, EDTA-free [Thermo Scientific]) and lysed. The cell resuspension was lysed in 4 mL batches. Lysis involved 10 rounds of sonication using a Fisher Scientific Sonic Dismembrator sonicator (FB-705) set at amplitude 10 for 10 s pulsing on and off for 1 s each, followed by a 20 s period between each sonication event. Thereafter, the resulting lysate was clarified by centrifugation at 15,000 *xg* for 15 min at 4 °C. Next, clarified lysate was applied to rabbit IgG agarose resin (5 mL, Sigma) pre-equilibrated with lysis buffer and incubated with the resin for 1 h at 4 °C on an end-over-end rocker. Unbound proteins were allowed to flow off the column by gravity. Subsequently, the resin bed was washed with 10 column volumes (CV) of lysis buffer without protease inhibitors and then with 5 CV of wash 2/cleavage buffer (25 mM HEPES, pH 7.4; 150 mL NaCl). 250 μL PreScission Protease was diluted in 5 mL cleavage buffer and used to resuspend the resin with the column capped. The column was incubated at 4 °C overnight without agitation.

The next morning, the cleaved protein was allowed to flow off the column by gravity and the column was washed two times with 5 CV of wash 2 buffer. These washes were combined with the cleaved protein, concentrated to < 500 µL with an Amicon Centrifugal Filter Unit MWCO 10,000, and applied to a 10/300 Supderdex 200 pg gel filtration column. Protein was eluted with 36 mL of wash 2 buffer. Protein purify was confirmed by SDS-PAGE and Coomassie staining. After confirming that the protein was >95% pure, fractions containing SARM1^ΔMLS^ were pooled and the protein was concentrated using an Amicon Centrifugal Filter Unit MWCO 10,000, aliquoted in 25 μL, flash frozen in liquid nitrogen, and stored at -80 °C. Protein concentration was determined by the Bradford method before freezing.

### Expression and purification of the TIR domain from SARM1

The expression and purification of the TIR domain from SARM1 was performed as previously described.^32,57^ Briefly, a pET-30a(+) vector expressing a Twin-Strep tag and the TIR domain from SARM1 (residues 561-724) were transformed into chemically competent *Escherichia coli* C43 (DE3), plated on LB agar supplemented with 50 µg/mL kanamycin, and incubated overnight at 37 °C. Single colonies were selected and cultured overnight at 37 °C with agitation in 5 mL LB with 50 µg/mL kanamycin. The following day, 2 L of LB plus kanamycin (50 µg/mL, final) were inoculated with the starter cultures and incubated at 37 °C with agitation until an OD_600_ = 0.8 was reached. The cultures were cooled on ice for 30 min and then expression was induced with 500 µM (final concentration) of IPTG. The TIR domain was expressed at 16 °C with agitation for 16-20 h. Cells were harvested at 5,000 *x g* and the pellets were flash frozen in liquid nitrogen before storing at -80 °C until use.

To purify the TIR domain, cell pellets from 10 L growths were thawed on ice and resuspended in 200 mL lysis buffer (100 mM HEPES [pH 8.0]; 200 mM NaCl; 10% glycerol [w/v]; 0.01% Tween20) supplemented with Pierce Protease Inhibitor Tablets, EDTA-free (Thermo Scientific). The resuspension was incubated with lysozyme (100 µg/mL, final concentration) for 10 min on ice. Cells were lysed by 12 rounds of sonication using a Fisher Scientific Sonic Dismembrator sonicator (FB-705) set at amplitude 30 for 20 s pulsing on and off for 1 s each, followed by a 30 s period between each sonication event. Following lysis, the crude lysate was clarified by centrifugation at 15,000 *x g* for 30 min at 4 °C. During clarification, 6 mL of Strep-Tactin XT 4Flow high-capacity resin (3 mL column volume [CV], IBA) was equilibrated with wash buffer (50 mM HEPES [pH 8.0]; 500 mM NaCl). The clarified lysate was flowed over the resin by gravity and then the resin was washed with 30 CV wash buffer. Thereafter, the protein was eluted with 20 CV elution buffer (50 mM HEPES [pH 8.0]; 500 mM NaCl; 50 mM biotin) and then concentrated to <2 mL using an Amicon Centrifugal Filter Unit MWCO 10,000. Concentrated protein was injected on a 16/600 HiLoad Superdex 200 pg gel filtration column and separated with 180 mL gel filtration buffer (50 mM HEPES [pH 8.0]; 150 mM NaCl). Protein purity was evaluated by SDS-PAGE and fractions containing the TIR domain from SARM1 were pooled, concentrated, and aliquoted to 25-100 µL volumes. The protein concentration was determined by the Bradford assay (BioRad), and then the aliquots were flash frozen in liquid nitrogen and stored at -80 °C until use.

### Evaluating SARM1^ΔMLS^ ribosylation in Expi239F lysates

To evaluate global ribosylation in response to SARM1 expression, Western blots were performed on lysates from Expi293F cells. Untransfected cells were used as a control, whereas SARM1^ΔMLS^ expressing cells were used for the experimental condition. Total protein (20 μg/well) was run on an SDS-PAGE gel and transferred to a PVDF membrane for 1 h at 80 V. The membrane was blocked in 5% bovine serum albumin (BSA) in phosphate buffered saline with 0.1% Tween- 20 (PBST) for 1 h at room temperature, after which the primary antibodies were added. The anti- MARylation antibody (Biorad, rabbit) was used at a 1:500 dilution and the anti-PARylation (Abcam, mouse) antibody was used at a 1:1000 dilution. Membranes were incubated with the primary antibody overnight at 4 °C. The next day, PBST was used to briefly wash the membrane and the secondary antibodies, diluted 1:5000 in 5% BSA in PBST, were incubated with the membrane for 1 h at room temperature in the dark. Two secondary antibodies were used: goat-anti-rabbit conjugated to IRDye680 (Licor) and goat-anti-mouse conjugated to IRDye800 (Licor). Membranes were washed twice with PBST and once with water. A Licor scanner equipped with Odyssey imaging software was used to image the blots.

### SARM1^ΔMLS^ mutants

The SARM1^ΔMLS^ mutants G601P, E642A, H685A, E189A, D594A, D632A, D634A, and E189A/D634A were generated by standard PCR-based site directed mutagenesis protocols. The pcDNA3.4(+) vector encoding SARM1^ΔMLS^ (residues 28-724) fused to a PreScission Protease site and two tandem Protein A tags was used as a template (0.2 ng/μL; final concentration). The manufacturer’s protocol for NEBNext High-Fidelity 2x PCR Master Mix was used without modification. 50 μL reactions were used in the following amplification conditions: initial denaturation for 30 s at 98 °C, denaturation for 10 s at 98 °C, annealing for 30 s at 50°C–72°C, extension for 4:40 min at 72 °C, final extension for 2 min at 72 °C. Denaturation, annealing, and extension steps were repeated 30 times. Next, to digest the template DNA, 2 μL DpnI (NEB) was added to the reactions and incubated at 37 °C for 2 h. The digest was subsequently transformed into XL1-Blue cells (Agilent). The next day, single colonies were cultured at 37 °C with agitation in 8 mL LB media supplemented with 100 μg/mL ampicillin and 10 μg/mL tetracycline•HCl. Following an overnight growth period, the plasmid was purified with a mini-prep kit (Promega). Mutagenesis was confirmed by Sanger sequencing (Genewiz) and the mutants were expressed and purified as for wild-type SARM1^ΔMLS^.

### Evaluating the ribosylation state of purified SARM1^ΔMLS^

Wild type, G601P, E642A, H685A, E189A, D594A, D632A, D634A, or E189A/D634A SARM1^ΔMLS^ (1.5 or 0.75 μg) was incubated with and without 1 mM NAD^+^ in triplicate for 15 min. The reaction was quenched with 5x SDS gel loading buffer and MAR/PARylation was analyzed by Western blot using the same conditions as above. MARylation was analyzed first. The anti- MARylation primary antibody (rabbit) was used at a 1:500 dilution and the IRDye680-conjugated goat-anti-rabbit antibody was used at a 1:5000 dilution. The blot was then stripped and reprobed for PARylation (if applicable) and SARM1 in parallel. The primary antibodies (mouse-anti- PARylation and rabbit-anti-SARM1 [CST], respectively) were used at a 1:1000 dilution and the secondary antibodies (IRDye800-conjugated goat-anti-mouse and IRDye680-conjugated goat-anti-mouse, respectively) were used at a 1:5000 dilution. After washing, the blots were imaged using a Licor scanner. The densities of the bands were analyzed in Image J and the MARylation or PARylation signals were normalized to the SARM1 signal. Densitometry values were further normalized to those for wild-type SARM1^ΔMLS^ in the absence of NAD^+^. Relative densities were plotted in GraphPad Prism.

### ENAD hydrolysis assay

The EADPR standard curve was generated as previously described.^31,33,57^ In brief, excess ADPR cyclase (Sigma) was incubated with 400 µM ENAD in 120 µL of 20 mM HEPES (pH 7.5) and NaCl in duplicate. Fluorescence intensity was monitored at 25 °C using λ_ex_ = 340 nm and λ_em_ = 405 every 60 s until a plateau was reached, approximately 30 min; the plate reader used was a PerkinElmer EnVision 2104 Multilabel Reader and Wallac EnVision Manager software. Thereafter, the reaction was serially diluted 1:2 six times and the final well was a buffer blank. Fluorescence intensity of the dilution series was measured once. Assuming all the ENAD reacted, the fluorescence intensity values were plotted against [EADPR] to yield the standard curve.

The ENAD assay was initiated with the addition of ENAD and monitored for 20 min, every 15 s, at 25 °C using λ_ex_ = 340 nm and λ_em_ = 405. Fluorescence intensities were converted to [EADPR] using the standard curve. Please see the descriptions below for experiments for information about concentrations, additives, and subsequent processing.

### The base exchange assay with NAD^+^ and PC6

The PAD6 standard curve was generated as previously described.^30,31^ In brief, excess TIR domain was incubated with 400 µM PC6 and 2000 µM NAD in 120 µL of 20 mM HEPES (pH 7.5) and NaCl in duplicate. Fluorescence intensity was monitored at 25 °C using λ_ex_ = 390 nm and λ_em_ = 420 every 60 s until a plateau was reached in approximately 2 h. The plate reader used was a PerkinElmer EnVision 2104 Multilabel Reader with monochromator functionality and Wallac EnVision Manager software. Thereafter, the reaction was serially diluted 1:2 six times and the final well was a buffer blank. Fluorescence intensity of the dilution series was measured once at 25 °C. Assuming the reaction went to completion, the fluorescence intensity values were plotted against [PAD6] to yield the standard curve.

The base exchange assay with PC6 was initiated with the addition of NAD^+^ and monitored for 20 min, every 15 s, at 25 °C using λ_ex_ = 390 nm and λ_em_ = 520. Fluorescence intensities were converted to [PAD6] using the standard curve. Please see the descriptions for experiments for information about concentrations, additives, and subsequent processing.

### Kinetic analysis of SARM1^ΔMLS^ mutants

In triplicate, wild type, G601P, E642A, H685A, E189A, D594A, D632A, D634A, or E189A/D634A SARM1^ΔMLS^ (750 nM) was combined with and without 25% PEG 3350 and 500 µM NMN in 20 mM HEPES (pH 7.5) and 150 mM NaCl (final concentrations). E189A, D594A, D632A, D634A, or E189A/D634A SARM1^ΔMLS^ were also evaluated with 25% PEG 3350 or 500 µM NMN alone. The mixture was incubated at room temperature for 10 min. ENAD hydrolysis was initiated by the addition of 0-1000 µM ENAD. The reaction was monitored for 20 min, every 15 s, at 25 °C using λ_ex_ = 340 nm and λ_em_ = 405. Using the EADPR standard curve (*vide supra*), fluorescence intensities were converted to [EADPR]. Base exchange was initiated with the addition of 0-1000 µM NAD^+^. The reaction was monitored for 20 min, every 15 s, at 25 °C using λ_ex_ = 390 nm and λ_em_ = 520. Using the PAD6 standard curve (*vide supra*), fluorescence intensities were converted to [PAD6]. The resulting progress curves were linear with respect to time and up to 10% turnover was used in subsequent analysis. Velocities were taken as the slopes of these lines and plotted in GraphPad Prism. Data were fitted to the Michaelis-Menten equation (**Equation 1**).

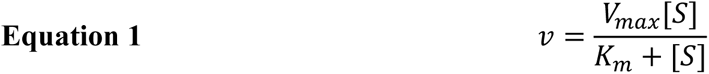

where *V_max_* is the maximum velocity, [S] is the substrate concentration, and *K_m_* concentration at half the maximum velocity.

### The Protein A affinity tag is PARylated

1.5 µg purified SARM1^ΔMLS^ or E642A SARM1^ΔMLS^ was added to 50 µL untransfected Expi293F lysates at 2 µg/µL in duplicate. Samples were incubated for 30 min at room temperature with and without 1 mM NAD^+^. Untransfected Expi293F lysates, pure protein, and lysates from Expi293F cells overexpressing Protein A-tagged wild type or E642A SARM1^ΔMLS^, with and without 1 mM NAD^+^, served as controls. Reactions were quenched with 1x SDS loading buffer. All samples (20 µg total protein) were analyzed by Western blotting against MARylation (BioRad), PARylation (Abcam), and SARM1 (CST). All primary antibodies were used at a 1:1000 dilution and secondary antibodies (IRDye800 goat-anti-rabbit and IRDye680 goat-anti-mouse, Licor) were used at a 1:5000 dilution. Blots were imaged on a Licor scanner with Odyssey software.

### Kinetics of the MARylation reaction in vitro

SARM1^ΔMLS^ (2.5 µM) was MARylated with 1 mM NAD^+^ for 30 min in a final volume of 80 µL assay buffer (20 mM HEPES, pH 7.5 and 150 mM NaCl) followed by 1:2 serial dilutions and the reaction was quenched with 10 µL 5X SDS loading buffer. 10µL of each reaction was loaded onto an SDS-PAGE gel and analyzed by Western blotting against MARylation. Band intensities were analyzed using ImageJ to generate a standard of [ADPR] produced. SARM1^ΔMLS^ (750 nM) was then mixed in 20 mM HEPES (pH 7.5), 150 mM NaCl with or without 500 µM NMN in triplicate. The MARylation reaction was initiated with 0-1000 µM NAD^+^ and incubated for 1 min at room temperature. Samples were quenched with SDS loading buffer and analyzed by Western blotting against MARylation. Band intensities were converted to [ADPR] using the standard curve described above and were analyzed in ImageJ and quantified with GraphPad Prism. The resulting values were fit to **Equation 1** for kinetic analysis.

### Inhibition of the MARylation reaction with known SARM1 inhibitors

SARM1^ΔMLS^ (750 nM) was incubated in 20 mM HEPES (pH 7.5), 150 mM NaCl with 0- 10 µM ZnCl_2_ or compound **10** for 30 min in duplicate. The MARylation reaction was initiated with 100 µM NAD^+^ and incubated for 5 min at room temperature. Samples were quenched with 10 µL 1x SDS loading buffer and analyzed by Western blotting against MARylation and SARM1. Images were analyzed in ImageJ and quantified with GraphPad Prism. MARylation densities were normalized to SARM1 densities and the 0 µM inhibitor condition was set at 100% activity. These normalized values were fitted to **Equation 2** for kinetic analysis,

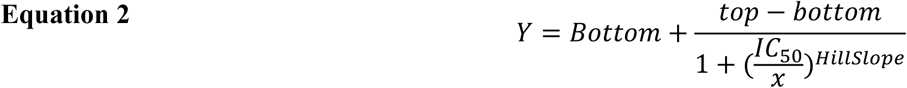

where top and bottom are upper and lower plateaus, respectively, IC_50_ is the concentration of inhibitor that gives a response halfway between top and bottom, and Hill Slope describes the steepness of the curve.

For inhibition of the ENAD hydrolysis reaction, SARM1^ΔMLS^ (750 nM) was incubated in 20 mM HEPES (pH 7.5), 150 mM NaCl with 0-10 µM ZnCl_2_ or compound **10** for 30 min in triplicate. The hydrolysis reaction was initiated with 100 µM ENAD and the reaction was monitored for 20 min, every 15 s, at 25 °C using λ_ex_ = 340 nm and λ_em_ = 405 on an Envision plate reader. Fluorescence intensities were converted to [EADPR] using the standard curve described above. Up to 10% of substrate turnover was analyzed further and the slopes of these lines were taken as the reaction velocity. The velocity for 0 µM inhibitor was set to 100% activity. Normalized data was fit to **Equation 2**.

### Hydroxylamine removes MARylation from SARM1^ΔMLS^

In triplicate, SARM1^ΔMLS^ (750 nM) was incubated in 20 mM HEPES (pH 7.5), 150 mM NaCl with 500 μM NAD^+^ for 30 min at room temperature. Next, 500 mM hydroxylamine (NH_2_OH) or water was added to the samples and incubated at 37 °C for 1 h. Reactions were quenched with 5x SDS gel loading buffer and analyzed by Western blot. Here, proteins were transferred at 100 V for 90 min. Membranes were blocked in 5% non-fat milk in PBST for 1 h at room temperature. All primary antibodies (rabbit-anti-MARylation and rabbit-anti-SARM1) were used at a 1:1000 dilution in 5% milk in PBST, and the secondary antibody was used at a 1:2500 dilution in 5% milk in PBST. In this experiment, the secondary antibody was a horseradish peroxidase (HRP)-conjugated goat-anti-rabbit antibody (Thermo). Chemiluminescent signal was developed with Radiance ECL (Azure Biosystems) for 5 min and detected on an Amersham imager.

### MacroD1 removes MARylation from SARM1ΔMLS

In duplicate, SARM1^ΔMLS^ (5 µM) was incubated in 20 mM HEPES (pH 7.5), 150 mM NaCl with or without 1000 µM NAD^+^ for 5 min at room temperature. Next, MARylated SARM1 was diluted to a final concentration of 750 nM in a reaction mix containing equimolar Macrodomain 1 (MacroD1) or water and the samples were incubated at room temperature for up to 1 h. At the specified time points, aliquots from the master mix were removed. Reactions were quenched with 5x SDS gel loading buffer and analyzed by Western blot against MARylation (BioRad) and SARM1 (CST). Here, proteins were transferred at 80 V for 60 min and the immunofluorescent Licor system was used for detection.

### MacroD1 dose response

SARM1^ΔMLS^ (750 nM) was incubated in 20 mM HEPES (pH 7.5), 150 mM NaCl with 0- 1000 nM MacroD1 for 5 min at room temperature in triplicate. After incubation with MacroD1, 500 µM PC6 was included and incubated for 10 min further for the base exchange reaction. The hydrolysis and base exchange reactions were initiated with 100 µM ENAD or 1 mM NAD^+^, respectively. The reactions were monitored for 20 min, every 15 s, at 25 °C using λ_ex_ = 340 nm and λ_em_ = 405 for the hydrolysis reaction or λ_ex_ = 390 nm and λ_em_ = 520 for the base exchange reaction on an Envision plate reader. Fluorescence intensities were converted to [EADPR] or [PAD6] using the standard curves described above. Up to 10% of substrate turnover was analyzed further and the slopes of these lines were taken as the reaction velocity. Velocities were plotted in GraphPad Prism.

### MacroD1 and hydroxylamine effect on SARM1^ΔMLS^ activity

SARM1^ΔMLS^ (750 nM) was incubated in 20 mM HEPES (pH 7.5), 150 mM NaCl with or without 250 nM MacroD1 or 500 mM neutral NH_2_OH for 1 h in at least triplicate. The no treatment control and MacroD1 samples were incubated at room temperature, whereas the NH_2_OH samples were incubated at 37 °C. The ENAD hydrolysis reaction was initiated with 0-1000 µM ENAD and the reaction was monitored for 20 min, every 15 s, at 25 °C using λ_ex_ = 340 nm and λ_em_ = 405 on an Envision plate reader. Fluorescence intensities were converted to [EADPR] using the standard curve described above. Up to 10% of substrate turnover was analyzed further and the slopes of these lines were taken as the reaction velocity. These data were fitted to **Equation 1** for kinetic analysis.

SARM1^ΔMLS^ (750 nM) was incubated in 20 mM HEPES (pH 7.5), 150 mM NaCl with or without 250 nM MacroD1 for 15 min at room temperature in triplicate. After this initial incubation, 25% PEG 3550, 500 µM NMN, and/or equal volumes of water were added to the mixtures and incubated for another 10 min. MacroD1 alone was used as a control. The hydrolysis reaction was initiated with 100 µM ENAD and the reaction was monitored for 20 min, every 15 s, at 25 °C using λ_ex_ = 340 nm and λ_em_ = 405 on an Envision plate reader. Fluorescence intensities were converted to [EADPR] using the standard curve described above. Up to 10% of substrate turnover was analyzed further and the slopes of these lines were taken as the reaction velocity. SARM1 in buffer was set to 100% activity and the normalized velocities were plotted in GraphPad Prism and analyzed by one-way ANOVA.

### MacroD1 effect on the phase transition

For SARM1^ΔMLS^, 2.5 µg protein was incubated in 20 mM HEPES (pH 7.5), 150 mM NaCl with 0, 0.25, or 1 µM MacroD1 for 30 min at room temperature in duplicate. For the TIR domain of SARM1, 5 µg protein was incubated in 20 mM HEPES (pH 7.5), 150 mM NaCl with or without 1 µM MacroD1 for 30 min at room temperature in duplicate. Next, 0-25% PEG 3350 was added, and the mixtures were incubated at room temperature for another 15 min. Samples were centrifuged at 21,000 *x g* for 10 min at room temperature, after which the soluble and insoluble fractions were separated. Protein was isolated by vortexing with 4 volumes of ice-cold acetone and storing at -20 °C for 1 h. Next, samples were brought to room temperature and centrifuged at 15,000 *x g* for 10 min at room temperature and air dried. Samples were resuspended in 10 µL 1x SDS loading buffer and analyzed by SDS-PAGE and Coomassie staining.

### MacroD1 effect on the NMN response of SARM1^ΔMLS^

SARM1^ΔMLS^ (750 nM) was incubated in 20 mM HEPES (pH 7.5), 150 mM NaCl with and without 250 nM MacroD1 at room temperature in triplicate. Next, 0-800 µM NMN was added, and the mixture was incubated for 5 min further. 500 µM PC6 was included in the second incubation for the base exchange reaction. The hydrolysis and base exchange reactions were initiated with 100 µM ENAD or 1 mM NAD^+^, respectively. The reactions were monitored for 20 min, every 15 s, at 25 °C using λ_ex_ = 340 nm and λ_em_ = 405 for the hydrolysis reaction or λ_ex_ = 390 nm and λ_em_ = 520 for the base exchange reaction on an Envision plate reader. Fluorescence intensities were converted to [EADPR] or [PAD6] using the standard curves described above. Up to 10% of substrate turnover was analyzed further and the slopes of these lines were taken as the reaction velocity. Velocities were plotted in GraphPad Prism and fitted to **Equation 3**,

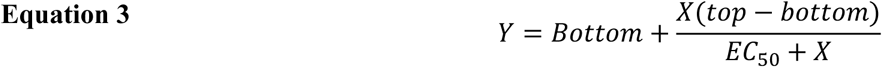

where top and bottom are the upper and lower plateaus, respectively, and the EC_50_ is the concentration of agonist that gives a response hallway between top and bottom.

### Determination of the sites of modification with mass spectrometry

SARM1^ΔMLS^ (0.5 mg/mL, 100 μL) was incubated in 20 mM HEPES (pH 7.5), 150 mM NaCl with and without 500 μM NAD^+^ (final concentration) for 15 min at room temperature to induce MARylation. Hydroxylamine (500 mM, final concentration) was added, and the samples were incubated at 37 °C for 1 h. Samples were diluted 2-fold with 100 μL of 8 M urea in 50 mM Tris, pH 8.0 (4 M urea final concentration). Next, the protein was reduced with 15 mM DTT (final concentration) at 70 °C for 20 min and alkylated with 12.5 mM iodoacetamide (final concentration) at 37 °C for 30 min in the dark. The volume of the sample was diluted by half with 50 mM Tris, pH 8.0, after which the protein was digested with trypsin (0.01 μg/μL final concentration) and CaCl_2_ (1 mM final concentration) overnight with end-over-end agitation at 37 °C. The next day, the digest was quenched with 20 μL formic acid and desalted with Pierce C18 Spin Columns (Thermo). Peptide concentration was determined by the Pierce Quantitative Colorimetric Peptide Assay (Thermo). Finally, peptides were dried using a speed vacuum concentrator and stored at - 80 °C until analysis.

For mass spectrometry analysis, peptides were reconstituted in 20 μL of 5% acetonitrile with 0.1% formic acid. A 2.0 μL sample was injected in triplicate onto a NanoElute LC coupled with a Bruker TimsTOF PRO2 MS system using a 30 min gradient. Data were searched against SwissProt human proteome database using Fragpipe/MSFragger and loaded to Scaffold 5 for visualization. Hydroxamic acid was included as a variable modification and SARM1 was in the first position (i.e., most abundant).

### Phase transition of MARylation mutants

Wild type, D594A, or D632A SARM1^ΔMLS^ (1.0 µg, 50 µL) was incubated with or without 1 mM NAD^+^ for 15 min at room temperature in 20 mM HEPES (pH 7.5), 150 mM NaCl in triplicate. Next, 25% PEG 3350 was added, and the mixtures were incubated at room temperature for another 15 min. Samples were centrifuged at 21,000 *x g* for 10 min at room temperature, after which the soluble and insoluble fractions were separated. Protein was isolated by vortexing with 4 volumes of ice-cold acetone and storing at -20 °C for 1 h. Next, samples were brought to room temperature and centrifuged at 15,000 *x g* for 10 min at room temperature and air dried. Samples were resuspended in 10 µL 1x SDS loading buffer and analyzed by SDS-PAGE and Coomassie staining.

### Hydroxylamine inhibits TIR domain activity

The TIR domain from SARM1 (2.5 µM) was incubated with or without 500 mM hydroxylamine at 37 °C for 1 in triplicate. Following this initial incubation, 20 mM HEPES (pH 7.5), 150 mM NaCl, and 25% PEG 3350 or an equal volume of buffer was added and samples were incubated at room temperature for another 10 min. The ENAD hydrolysis reaction was initiated with 500 µM ENAD and the reaction was monitored for 20 min, every 15 s, at 25 °C using λ_ex_ = 340 nm and λ_em_ = 405 on an Envision plate reader. Fluorescence intensities were converted to [EADPR] using the standard curve described above. Up to 10% of substrate turnover was analyzed further and the slopes of these lines were taken as the reaction velocity. The TIR domain without hydroxylamine and with PEG 3350 was set to 100% activity. Normalized velocities were plotted in GraphPad Prism and analyzed by one-way ANOVA.

### SARM1 catalyzes MARylation in trans in vitro

E642A SARM1^ΔMLS^ (750 nM) was incubated in 20 mM HEPES (pH 7.5), 150 mM NaCl, 25% PEG 3350, 1 mM NAD^+^, and 0-4 µM TIR domain from SARM1 for 15 min in duplicate. Controls for this experiment include the TIR domain alone and E642A SARM1^ΔMLS^ alone, with and without 1 mM NAD^+^. The MARylation reaction was quenched by vortexing with 4 volumes of ice-cold acetone and storing at -20 °C for 1 h. Next, samples were brought to room temperature and centrifuged at 15,000 *xg* for 10 min at room temperature and air dried. Samples were resuspended in 10 µL 1x SDS loading buffer and analyzed by Western blotting against SARM1 (CST) and MARylation (BioRad). Proteins were transferred at 80 V for 60 min and the immunofluorescent Licor system was used for detection. The same experimental protocol was used for PAD2.

### SARM1 catalyzes MARylation in cells

To study MARylation catalyzed by SARM1 in cells, tag-free or Protein A-tagged SARM1^ΔMLS^ were transfected into Expi293F cells using the methods described above in triplicate. Cells from 10 mL cultures were harvested by centrifugation and cell pellets were flash frozen and store at -80 °C until use. Untransfected Expi293F cells were used as a control.

To study the MARylation activity of endogenous SARM1, 11 x 10^6^ SHSY5Y cells were seeded in 100 mm dishes in DMEM:F12 supplemented with 10% fetal bovine serum (FBS, Gibco) and 1% penicillin/streptomycin (Gibco) and allowed to adhere overnight at 37 °C and >80% relative humidity. The following morning, cells were washed once with 1x PBS and then the media was switched to serum free DMEM:F12 with 1% penicillin/streptomycin. In triplicate, cells were treated for 2 h with 300 µM 5-iodoisoqinoline or vacor in DMSO and DMSO was used as the control; the final DMSO concentration was 0.3%. After 2 h, the media was discarded, and the cells were harvested by cell scraping in 10 mL 1x PBS.

For both experiments, cells were lysed by subcellular fractionation.^61,62^ Briefly, cells were resuspended in 750 µL Hypotonic buffer (10 mM Tris·HCl, pH 7.5; 10 mM NaCl; 1.5 mM MgCl_2_) and incubated on ice for 5 min. The resuspension was transferred to a 2 mL Dounce homogenizer and cells were lysed with 40-50 strokes of the B pestle. 500 µL 2.5x Homogenization buffer (12.5 mM Tris·HCl, pH 7.5; 525 mM mannitol; 175 mM sucrose; 2.5 mM EDTA) was triturated with the lysed cells. Lysate was transferred to a 2 mL microcentrifuge tube and the douncer was washed with 625 µL 1x Homogenization buffer (5 mM Tris·HCl, pH 7.5; 210 mM mannitol; 70 mM sucrose; 1 mM EDTA). The wash was added to the lysate. Next, samples were centrifuged at 700 *x g* for 10 min at 4 °C to remove unlysed cells, nuclei, and large fragments of cell membranes. The supernatant was poured into another microcentrifuge tube and centrifuged again at 12,000 *x g* for 15 min at 4 °C to pellet the mitochondria. The supernatant from this step was retained as the cytosolic fraction and dried in a vacuum concentrator. Mitochondrial pellets were stored at 4 °C overnight.

The following day, the cytosolic proteins were reconstituted in 500 mL RIPA buffer, whereas mitochondria were lysed in 100 µL RIPA buffer. Protein concentration was determined by the Detergent Compatible Protein Assay (BioRad) and 40 µg of total protein from each sample was analyzed by Western blotting against SARM1 (CST), MARylation (BioRad), and CoxIV (Thermo). Proteins were transferred to PVDF membrane at 100 V for 90 min and the membrane was blocked in 5 % milk in PBST for 1 h. Primary antibodies were used at 1:1000 dilution in 1% milk in PBST and HRP-secondary antibodies were used at 1:3000 dilution in 2.5% milk in PBST. SuperSignal West Femto Maximum Sensitivity Substrate (Thermo) and an Amersham imager were used for detection.

## Data, Materials, and Software Availability

All source data is available from the authors upon request. All study data are included in the article and/or SI Appendix.

## Acknowledgement and Funding Sources

This work was supported in part by National Institutes of Health grants <u>R35 GM118112</u> (P.R.T.), <u>T32 AI132152</u> (L.D.), and <u>F31 NS122423</u> (J.D.I.), and the Dan and Diane Riccio Fund for Neuroscience (P.R.T.). The content is solely the responsibility of the authors and does not necessarily represent the official views of the National Institutes of Health.

## Author contributions

JDI, LD, and PRT designed research; JDI and LD performed experiments; LB synthesized 10; JDI and LD analyzed experiments; BAK and PRT provided experimental advice and supervised the research; JDI and PRT wrote and edited the paper.

