## Supporting information for "SARM1, the executioner of axon degeneration, is an ADP-ribosyl transferase and autoMARylation negatively regulates its activation"

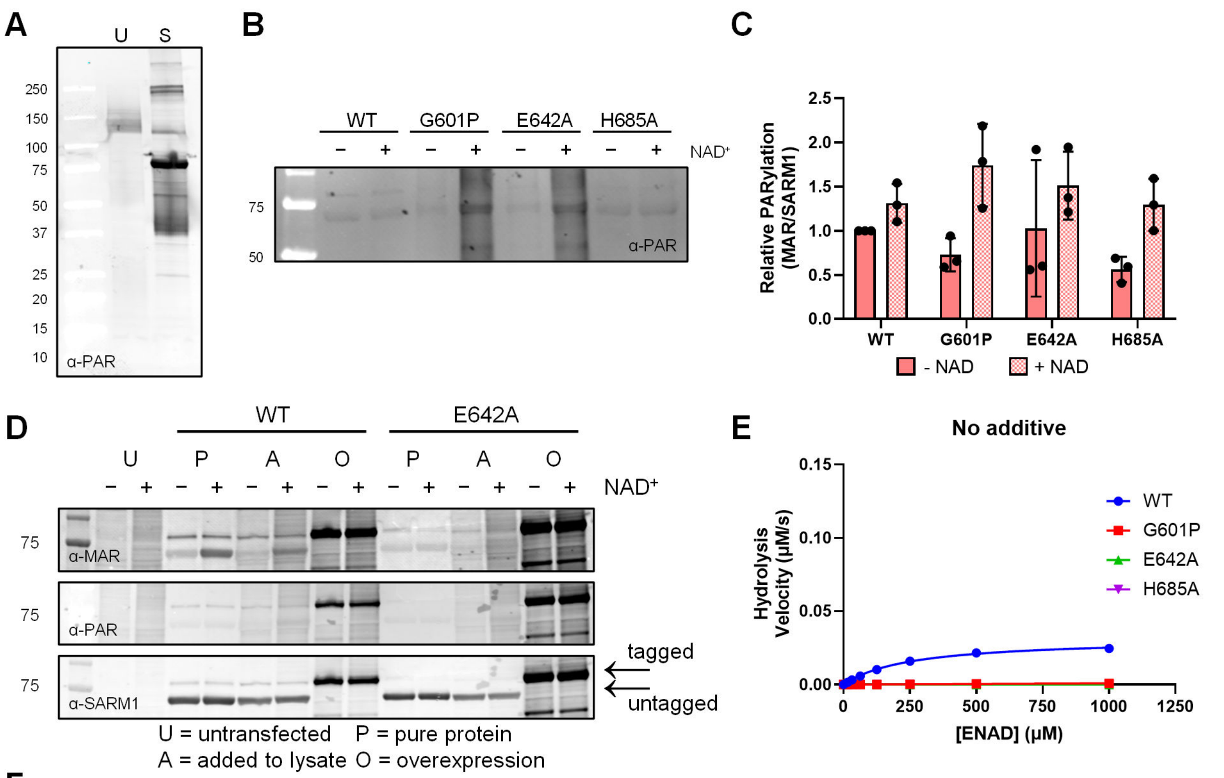


F

| | Mutant | No additive $K_m$ (µM) | No additive $k_{cat}$ ($s^{-1}$) | No additive $k_{cat}/K_m$ ($M^{-1}s^{-1}$) | 25% PEG 3350/500 µM NMN $K_m$ (µM) | 25% PEG 3350/500 µM NMN $k_{cat}$ ($s^{-1}$) | 25% PEG 3350/500 µM NMN $k_{cat}/K_m$ ($M^{-1}s^{-1}$) |
|---|---|---|---|---|---|---|---|
| Hydrolysis | WT* | 250 ± 16 | 0.042 ± 0.001 | 166 ± 5 | 180 ± 8 | 0.21 ± 0.003 | 1200 ± 7 |
|  | G601P | ---- | ---- | 1.32 | ---- | ---- | ---- |
|  | E642A | ---- | ---- | ---- | ---- | ---- | ---- |
|  | H685A | ---- | ---- | ---- | 23 ± 18 | 0.004 ± 0.001 | 160 ± 100 |

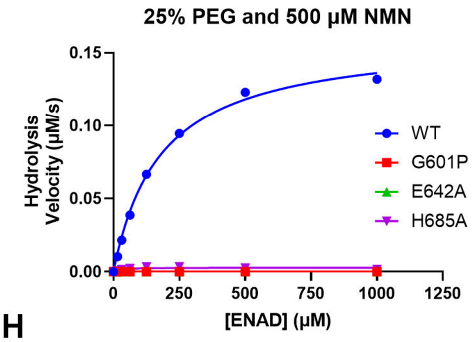


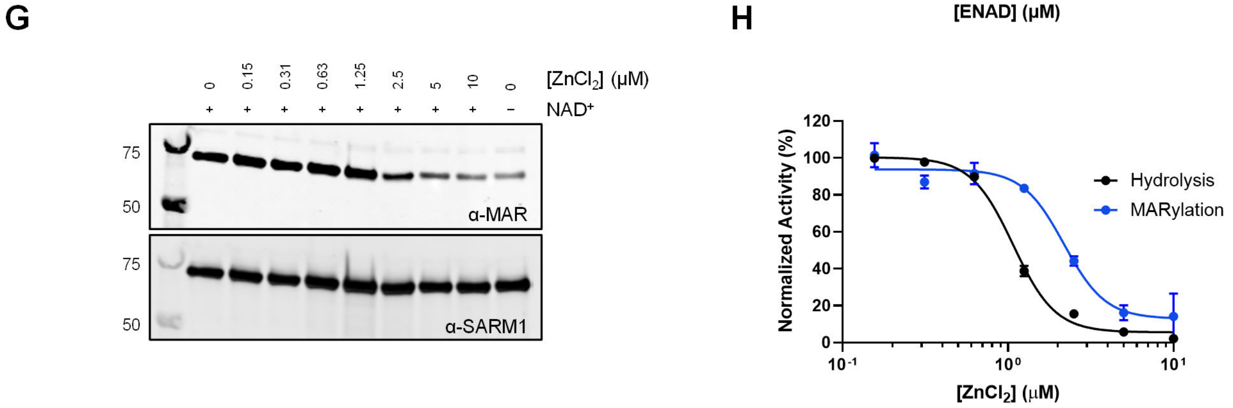


**Figure S1.** A) Recombinantly expressed SARM1 is poly-ADP-ribosylated in Expi293F lysates. U: untransfected; S: SARM1$^{\Delta MLS}$ overexpression. n = 3, representative image shown. B) Purified recombinant SARM1 does not have PARylation activity. Purified WT or variant SARM1$^{\Delta MLS}$ incubated with or without 1 mM NAD$^+$ for 30 min and analyzed by Western blot. n = 3, representative image shown. C) Quantification of B. Error reported as SD. D) SARM1$^{\Delta MLS}$

catalyzes MARylation, but not PARylation. U = untransfected Expi293F lysate; n = 2. E) Top – Kinetic analysis for the ENAD hydrolysis reaction of SARM1$^{\Delta MLS}$ variants in buffer. Bottom – Kinetic analysis for the ENAD hydrolysis reaction of SARM1$^{\Delta MLS}$ variants in the presence of 25% PEG 3350 and 500 μM NMN. n = 3, error reported as SEM. F) Summary of the kinetic parameters for SARM1$^{\Delta MLS}$ mutants for the hydrolysis reaction; n = 3, error reported as SD. G) Inhibition of the MARylation reaction by the SARM1 inhibitor $ZnCl_2$. SARM1$^{\Delta MLS}$ was incubated with 0-10 μM $ZnCl_2$ and then with 1 mM $NAD^+$ and analyzed by Western blot. n = 3, representative image shown. H) Quantification of F. Error reported as SD. Note: in some cases, the error is smaller than the size of the data point and is not visible.

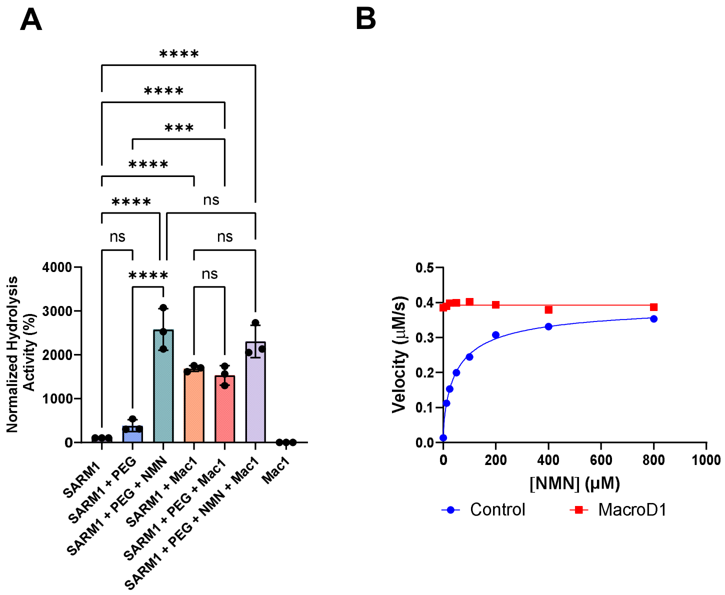


**Figure S2. MARylation regulates SARM1 activity.** A) ENAD hydrolysis activity of SARM1$^{\Delta MLS}$ incubated with and without 25% PEG 3350 and/or 500 µM NMN in the presence or absence of 250 nM MacroD1. n = 3, error reported as SD. B) NMN dose response in the base exchange reaction of SARM1$^{\Delta MLS}$ treated with and without 250 nM MacroD1; n = 3, error reported as SD. Note: in some cases, the error is smaller than the size of the data point and is not visible.

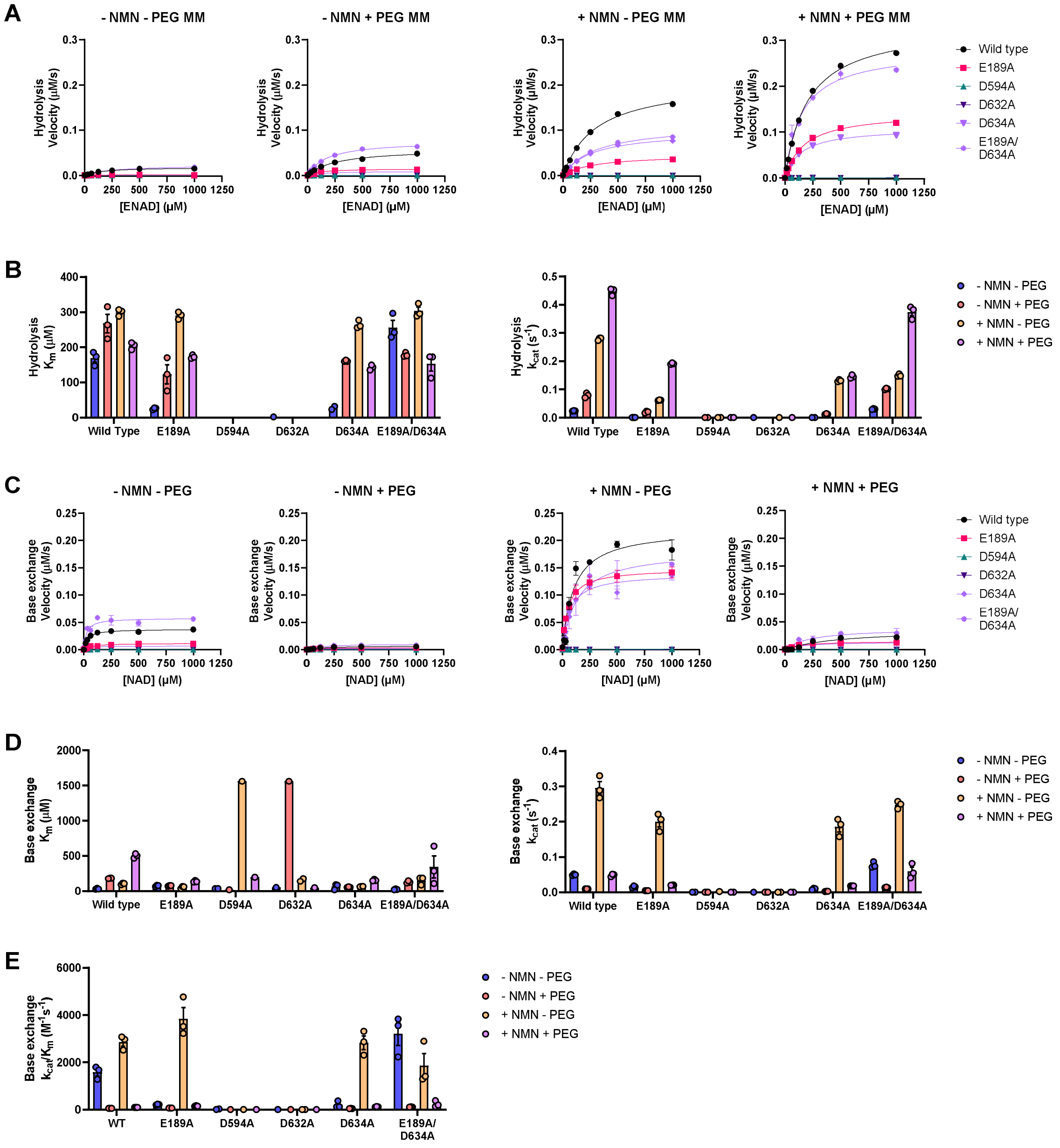


**Figure S3. Raw data associated with Figure 4.** A) Michaelis Menten curves for the hydrolysis reaction. B) Km and kcat for the hydrolysis reaction. C) Michaelis Menten curves for the base exchange reaction. D) Km and kcat for the base exchange reaction. E) Catalytic efficiency with respect to the base exchange reaction of MARylation variants treated with 0-1000 µM ENAD. n = 3, error reported as SD. Note: in some cases, the error is smaller than the size of the data point and is not visible.

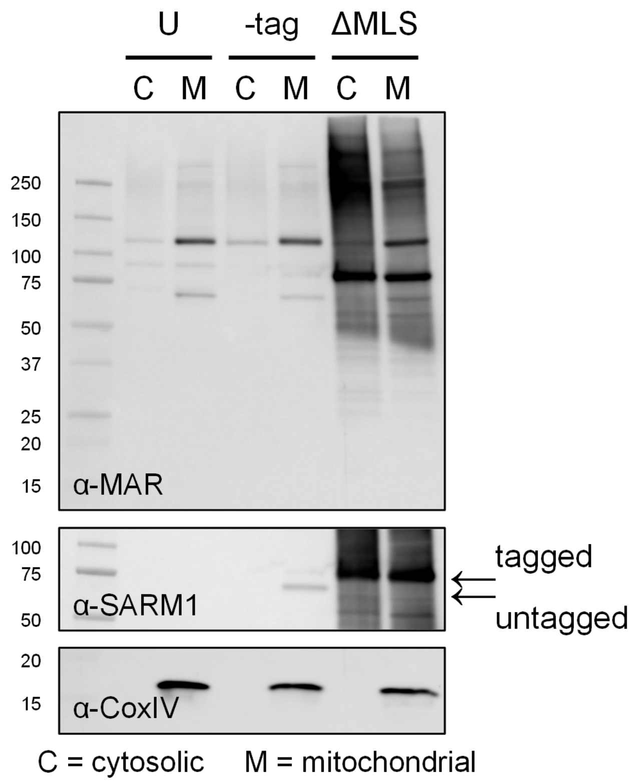


**Figure S4. SARM1 can catalyze MARylation in cells.** Western blot for SARM1, MARylation, and CoxIV from SHSY-5Y cells following overexpression of tag-free (-tag) or Protein A-tagged (ProA) SARM1$^{\Delta MLS}$ and subcellular fractionation (C = cytosolic, M = mitochondrial). n =3; representative images shown.
